# T cell intrinsic 4-1BB signals induce Prdm16 to increase effector and memory T cell numbers during respiratory influenza infection

**DOI:** 10.64898/2026.04.02.716118

**Authors:** Seungwoo Lee, Karen Yeung, Carolina de Amat Herbozo, Razieh Eshraghisamani, Alina Dorogy, Tania H. Watts

## Abstract

TNFR superfamily members such as 4-1BB sustain T cell responses to control virus infections or tumors. However, the precise role of 4-1BB during an acute infection remains incompletely understood. Here we used mixed bone marrow chimeras and transcriptome analysis to show that intrinsic 4-1BB signaling in lung T cells during influenza A virus (IAV) infection induces the transcriptional coregulator PR domain containing 16 (*Prdm16),* known for its role in regulating mitochondrial biology in other cell types. T cell-specific deletion of *Prdm16* reduced the number of Ag-specific CD8 T cells, with a larger effect on T cells in the lung parenchyma compared to the vasculature or lymphoid tissues. Conversely, *Prdm16* overexpression in T cells increased effector and memory CD8 T cell accumulation during IAV infection. Single nuclei transcriptomics suggested that *Prdm16* allows the accumulation of T cells with high protein translation and mitochondrial activity. *Prdm16* increased genes associated with oxidative phosphorylation and mitophagy. Consistently, *Prdm16* overexpressing cells had more compact mitochondrial cristae, which has been associated with more efficient electron transport*. Prdm16* also repressed some genes, including Herpes virus entry mediator, which can inhibit T cell responses through B and T lymphocyte attenuator. These findings reveal a 4-1BB-Prdm16 axis that is induced in T cells during viral infection to support T cell accumulation and memory formation.

## Introduction

T cells must be precisely regulated at each stage of their lifespan to control immunity while minimizing collateral damage. The ability of T cells to expand and become memory T cells requires the regulation of multiple processes, including proliferation, survival, differentiation, and metabolic reprogramming.^1^ Understanding the intrinsic factors that control T cell responses in specific contexts is critical to fully understand the regulation of immunity. During T cell priming, signals through the TCR, CD28, as well as cytokine-dependent signaling drive antigen (Ag)-specific T cells to undergo clonal expansion and differentiation.^2, 3^ Subsequently, costimulatory TNFR family members on activated T cells engage their ligands post-priming to sustain T cell responses, particularly in the tissues.^4, 5^

The TNFR superfamily member 4-1BB (also known as CD137 or *Tnfrsf9*) has been implicated in CD8 T cell accumulation and memory formation during viral infection and cancer.^5–11^ 4-1BB critically regulates survival of activated T cells through upregulation of prosurvival Bcl-2 family members and downregulation of the proapoptotic molecule BIM.^12–16^ Agonistic anti-4-1BB antibody- or chimeric antigen receptor (CAR)-induced 4-1BB signaling in T cells promotes mitochondrial biogenesis and fusion, thereby contributing to increased respiratory capacity of the activated T cells.^17–19^ In the tumor microenvironment, 4-1BB drives a Tox-dependent chromatin remodelling program which increases the accumulation of exhausted T cells that, despite their exhausted state, contribute to tumor control.^20^ In contrast, in CD19-CAR-T cells, the 4-1BB signaling domain drives an enriched central memory (Tcm) phenotype, reduced hypoxia and increased fatty acid metabolism.^21, 22^ Thus, the downstream consequences of 4-1BB signaling in T cells are context dependent. In this regard, the mechanisms by which endogenous 4-1BB signaling contributes to T cell accumulation in an acute infectious disease setting remains incompletely understood.

Here we investigated the T cell intrinsic role of 4-1BB during acute respiratory influenza A virus (IAV) infection using mixed bone marrow chimeras and single cell RNA-sequencing. We find that T cell intrinsic 4-1BB signaling drives the accumulation of multiple Ag-specific effector and memory precursor CD8 T cell subsets, with greater effects in the lung compared to the spleen and with no discernible effect on CD8 T cell differentiation. Bulk RNA sequencing showed that T cell intrinsic 4-1BB signaling in the lung induces the transcriptional and epigenetic coregulator PR domain containing 16 (*Prdm16*). Although not previously studied in αβ T cells, Prdm16 is known for its role as a master regulator of brown adipose tissue identity, through interacting with other transcription factors such as PGC-1α to increase their transcriptional activity.^23–27^ Prdm16 also maintains hematopoietic stem cells (HSC) with lymphoid potential.^28, 29^ In HSC, a short isoform of Prdm16 induces Mitofusin 2, responsible for mitochondrial fusion and tethering to the endoplasmic reticulum.^28^ Prdm16 also regulates oxidative stress in HSC and neural stem cells^30^ and in cardiac tissues deletion of *Prdm16* leads to mitochondrial dysfunction.^31^ Prdm16 also has epigenetic activities: it promotes the brown fat transition^32^ by suppressing white fat specific genes through recruitment of the histone methyltransferase Ehmt1,^26^ and has repressive H3K9 methyltransferase activity in mouse embryonic fibroblasts.^33^ However, to date, the role of Prdm16 in αβ T cells has not been investigated. Here we show that deletion of *Prdm16* in mature mouse T cells results in decreased Ag-specific effector and memory T cell accumulation during IAV infection, particularly in the lung tissue, with the largest effect on lung resident memory T cells (Trm). Conversely, overexpression of full length *Prdm16* in OT-I CD8 TCR transgenic cells increases Ag-specific T cell accumulation across all CD8 T cell subsets in the lung and secondary lymphoid organs. Bulk RNA-sequencing as well as sequencing of single nuclei, suggests a role for Prdm16 in T cells in regulating multiple processes including oxidative phosphorylation and lipid metabolism. Both single nuclei and bulk RNA-sequencing revealed *Prdm16*-dependent-induction of *Pink1,* involved in the Pink/Parkin/DJ-1 mitophagy pathway,^34, 35^ a mitochondrial quality control mechanism that contributes to T cell memory.^36^ Consistently, *Prdm16* overexpression resulted in more compact mitochondrial cristae compared to mock transduced cells, a state that has been associated with improved oxidative phosphorylation.^37, 38^ In addition, Prdm16 repressed expression of Herpes virus entry mediator (HVEM), which can act as a ligand for B and T lymphocyte attenuator (BTLA) on T cells to limit T cell activation.^39–41^ These data support a role for T cell intrinsic Prdm16 in promoting T cell accumulation and memory T cell persistence, potentially through improving mitochondrial fitness as well as through limiting HVEM-dependent T cell inhibition.

## Materials and Methods

### Mice

C57BL/6 mice were obtained from The Jackson Laboratory (Bar Harbor, ME, USA). Age-matched female mice were used in all experiments. C57BL/6 4-1BB^-/-^ mice were originally provided by B. Kwon (National Cancer Institute Goyang, Korea).^42^ Conditional *Prdm16* knockout mice (*Prdm16^fl/fl^ dLck-Cre^+/–^*, referred to as *Prdm16* KO) were generated by crossing B6.129-Prdm16tm1.Brsp/J mice^27^ (Jackson), which contain loxP sites flanking exon 9 of *Prdm16*, with B6.Cg-Tg(Lck-icre)3779Nik/J mice expressing Cre recombinase under the distal *Lck* promoter to achieve deletion in mature T cells.^43^ To generate a 50:50 mix of WT and KO littermate mice, we bred *Prdm16^fl/fl^ dLck-Cre^+/–^* with *Prdm16^fl/fl^ Cre^-/-^* mice. *Prdm16^fl/fl^ dLck-Cre^+/–^* mice were crossed to CD45.1 OT-I mice (provided by David Brooks, Princess Margaret Cancer Centre, Toronto, ON). For Supplemental Fig.2B, we used OT-I *Prdm16^fl/fl^ dLck-Cre^+/–^*and OT-I *Prdm16^fl/fl^*Cre^-/-^ mice. For *in vivo* adoptive transfer experiments (Figure 4), we generated OT-I CD45.1 *Prdm16^+/+^ dLck-Cre^+/–^* (OT-I *Prdm16* WT) and OT-I CD45.2 *Prdm16^fl/fl^ dLck-Cre^+/–^* (OT-I *Prdm16* KO) strains.

For bone marrow chimera experiments, Thy1.1 (B6.PL-Thy1a/CyJ) and CD45.1 (B6.SJL-PtprcaPepcb/BoyJ) mice were obtained from The Jackson Laboratory. For OT-I adoptive transfer experiments, CD45.1 and CD45.2 C57BL/6 mice were bred in-house to generate CD45.1/2 heterozygous recipients. All mice were maintained under specific pathogen-free conditions in the Division of Comparative Medicine at the Terrence Donnelly Centre for Cellular and Biomolecular Research, University of Toronto. All animal procedures were approved by the University of Toronto Animal Care Committee and performed in accordance with the guidelines of the Canadian Council on Animal Care.

### Influenza virus infection

Influenza A/PR8 H1N1 (PR8) was used for infection of littermate mice, whereas the milder recombinant strain, Influenza A X31 H3N2 (X31), was used for mixed bone marrow chimeras due to the increased sensitivity of the radiation chimeras to influenza. The X31 strain shares the same internal proteins/CD8 T cell epitopes as Influenza A/PR8.^44^ For experiments with OT-I T cells, we used Influenza PR8-OVA, originally obtained from Paul Thomas (St. Jude Children’s Research Hospital, Memphis, TN, USA). For all viral strains, Infectious titers (tissue culture infectious dose 50; TCID_50_) were quantified by Madin-Darby canine kidney (MDCK) cell assays.^45^ Mice were anesthetized with isoflurane and intranasally (i.n.) infected with 30 μL of virus at a dose of 5 HAU (3.86×10^4^ TCID_50_) for X31, or 2.5×10^5^ TCID_50_ for PR8 or PR8-OVA. Animals were monitored daily and euthanized upon predefined endpoints.

### Mixed Bone Marrow Chimeras

Thy1.1 mice were lethally irradiated with two doses of 550 cGy and reconstituted intravenously with a 1:1 mixture of Thy1.2 CD45.1 WT and either Thy1.2 CD45.2 *4-1bb^-/-^* or *Prdm16* KO bone marrow cells (5×10^6^ total cells per mouse). Chimeric mice received drinking water supplemented with neomycin sulfate (2 mg/mL; Bio-Shop, Burlington, ON, Canada) for 2 consecutive weeks and were rested for >90 days to allow hematopoietic reconstitution before assessing blood chimerism. Mice were subsequently infected according to the experimental schedule.

### T-cell isolation and adoptive transfers

OT-I cells were enriched from spleens of CD45.1 *Prdm16* WT and CD45.2 *Prdm16* KO OT-I mice using the EasySep Mouse CD8 T-cell Isolation Kit (StemCell Technologies, Vancouver, BC, Canada). Cells were counted by trypan blue exclusion in triplicate and mixed at a 1:1 ratio. The input ratio and purity of OT-I cells were confirmed by flow cytometry based on CD3, CD8, TCR Vα2, TCR Vβ5, CD45.1, and CD45.2 expression. Mixed cells were adoptively transferred intravenously (200μL per mouse) into recipient mice 1 day prior to infection at the indicated cell numbers.

### Tissue harvest and processing

To distinguish lung parenchymal as compared to vascular exposed T cells, mice were injected intravenously with 3μg of fluorochrome-conjugated anti-mouse Thy1.2 or anti-mouse CD45 antibody and euthanized 10 min later.^46^ Lungs were perfused with 10mL of PBS, minced, and digested with 2mg/mL collagenase IV (Invitrogen, Carlsbad, CA, USA) for 30 min at 37°C with agitation. Tissue was then mechanically dissociated through a 70μm cell strainer, followed by red blood cell (RBC) lysis. Spleens and mLN were mechanically disrupted through a 70μm cell strainer to generate single-cell suspensions, followed by RBC lysis. Blood was collected via the saphenous vein and treated with RBC lysis buffer.

### T cell analysis by flow cytometry

Cell suspensions prepared from lungs, mLN, and spleens were Fc-blocked with anti-mouse CD16/32 monoclonal antibodies (eBioscience) for 15 min at 4°C, then stained with surface marker antibodies and Live&Dead dye for 30 min at 4°C. When necessary, H-2D^b^–restricted tetramer staining was performed simultaneously with surface antibody staining. Cells were washed and fixed in 4% paraformaldehyde (PFA) before flow cytometry analysis.

To detect influenza-specific CD8 T cells, biotinylated H-2D^b^/NP_366–374_ (ASNENMETM) or H-2D^b^/PA_224–233_ (SSLENFRAYV) monomers were obtained from the National Institute of Health (NIH) Tetramer Core Facility (Emory University, Atlanta, GA, USA), and tetramerized with streptavidin-APC or streptavidin-PE (ProZyne, San Leandro, CA, USA). Tem were defined as CD44^high^, CD62L^low^, CD69^-^, whereas Trm were defined as CD69^+^ and subdivided into CD103^+^ or CD103^-^ Trm.

For intracellular staining, cells were fixed and permeabilized using the Transcription Factor Staining Buffer Set (eBioscience) following the manufacturer’s guidelines. Cells were then labelled with fluorochrome-conjugated antibodies for 30min at 4°C. For cytokine analyses, lung and spleen homogenates were restimulated *ex vivo* with 1μM SIINFEKL peptide (OT-I experiments, 6hr) or live PR8 virus (18hr) at 37 °C, with BD GolgiStop present for the final 6hr. After restimulation, cells were subjected to surface staining, followed by fixation and permeabilization using BD Cytofix/Cytoperm Fixation/Permeabilization Kit (BD Biosciences) for intracellular cytokine staining. Unstimulated samples served as baseline controls. To assess mitochondrial membrane potential, reactive oxygen species, and fatty acid uptake, freshly isolated cells were incubated with MitoTracker Deep Red (Invitrogen), TMRM (Invitrogen), MitoSOX Red (Invitrogen), or BODIPY-FL C16 (Invitrogen) for 30min at 37°C according to manufacturers’ guidelines.

Flow cytometric data were acquired on LSR Fortessa, LSR Fortessa X-20, FACSymphony A3, or FACSymphony A5 instruments (BD Biosciences). Data analysis was performed using FACSDiva and FlowJo (Tree Star) software. To account for staining variability between independent experiments to allow pooling into one graph, dMFIs (MFI minus the corresponding FMO) from one batch were normalized using a correction factor calculated as the ratio of mean control (wildtype or mock-transduced) MFIs between batches.

### Prdm16 Overexpression

MSCV plasmids encoding EGFP or Prdm16 were obtained from Addgene (#20672 and #15504, respectively; Watertown, MA, USA). To generate a *Prdm16*-overexpressing construct with EGFP as a transduction marker, EGFP was cloned into the MSCV-*Prdm16* plasmid under the PGK promoter at the Genome Editing and Molecular Biology (GEM) Facility, University of Ottawa (Ottawa, Canada). MSCV-EGFP was used as a mock-transduced control. Phoenix packaging cells were transfected with these plasmids using Lipofectamine 3000 (Invitrogen, Carlsbad, CA, USA) according to the manufacturer’s instructions. Viral supernatants were collected and applied to cell-culture plates coated with RetroNectin (Takara Bio, San Jose, CA, USA) following manufacturer protocols.

CD45.1 or CD45.2 WT OT-I cells were enriched from spleens of WT OT-I mice and activated with anti-CD3 and anti-CD28 antibodies for 16hr, followed by IL-2 treatment (80 U/mL; Gibco, Cranbury, NJ, USA) for 2hr. Activated OT-I cells were transduced on virus-coated RetroNectin plates in the presence of IL-2 (20 U/mL) and IL-7 (10 ng/mL; StemCell Technologies, Vancouver, BC, Canada). Transduced cells were cultured in IL-2 and IL-7 for 8 days prior to adoptive transfer, electron microscopy, or flow cytometric analyses. For adoptive transfer, electron microscopy, and selected flow cytometry experiments, transduced cells were sorted by flow cytometry based on EGFP expression.

### Electron Microscopy

Mock-transduced and *Prdm16*-overexpressing OT-I cells were fixed in 4% paraformaldehyde and 1% glutaraldehyde in 0.1M phosphate buffer (pH 7.2) for 1hr at room temperature. After washing three times in phosphate buffer, 1% osmium tetroxide was added to post-fixation samples for 1hr in the dark then washed again in buffer. Dehydration was performed through a graded ethanol series followed by propylene oxide, and samples were infiltrated with epoxy resin mixtures of increasing concentration before polymerization at 60°C for 48 h. Ultrathin sections (∼90 nm) were cut using a Reichert Ultracut E ultramicrotome, collected on 300-mesh copper grids, and stained with 5% aqueous uranyl acetate and Reynold’s lead citrate. Sections were examined using a FEI Talos L120C transmission electron microscope operated at 120 kV. Mitochondrial number per cell, mean area of mitochondria, and cristae thickness were quantified with the person analyzing blinded as to sample origin.

### Sample preparation and raw data processing for bulk RNA sequencing

For 4-1BB experiments, CD45.1^+^ (*4-1bb^+/+^*WT) and CD45.2^+^ (*4-1bb^-/-^* KO) CD44^high^ CD8 T cells were FACS-sorted from lungs of infected mixed bone marrow chimeric mice. For *Prdm16* studies, CD45.1^+^ (WT or Mock) or CD45.2^+^ (KO or *Prdm16* OE) CD44^high^ OT-I cells were FACS-sorted from CD45.1/2 recipient mice that had been adoptively transferred with OT-I cells and infected as described above. Sorted cells were treated with 500μL LRT buffer (Qiagen, Hilden, Germany) and stored at −80°C with 1% β-mercaptoethanol until RNA extraction with the RNeasy Plus Micro Kit (Qiagen) according to the manufacturer’s instructions. Samples from 4-1BB and *Prdm16* studies were sequenced on the Illumina NovaSeq 6000 and NovaSeq X platforms, respectively.

### Bulk RNA-seq analysis

The UMI count matrix was filtered to retain genes with more than eight counts present in at least four samples. Batch factor was incorporated into the design matrix for 4-1BB experiments to account for variability introduced by experimental batch. Differentially expressed genes (DEGs) between experimental groups were identified using edgeR^47^ (4-1BB experiments) or DESeq2^48^ (Prdm16 experiments). Heatmap plots were generated from regularized log- transformed normalized counts. Gene Set Enrichment Analysis (GSEA) was performed on normalized counts using GSEA v4.3.2 with the Gene Ontology (GO) Biological Processes database. Gene regulatory networks were evaluated using the decoupleR package.^49^

### Single-cell CITEseq sample preparation

Lung and spleens were harvested from 4-1BB mixed bone marrow chimeras 8 days after i.n. infection with influenza PR8 and processed into single cell suspensions, pooled from 3 mice per organ. Samples were stained separately with fluor-conjugated antibodies (for FACS sorting) and TotalSeq-C antibodies (for sample and genotype demultiplexing). Both samples were labelled with anti-CD45.1 and anti-CD45.2 barcoded antibodies to distinguish 4-1BB WT (CD45.1) and 4-1BB KO (CD45.2) donor cells, while lung and spleen samples were labelled with different hashtag antibodies. Thy1.1^-^CD3^+^CD8^+^CD44^high^ cells were FACS sorted and submitted to the Princess Margaret Genomics Centre for CITE-sequencing.^50^ Lung and spleen samples were pooled together at a 1:1 ratio directly before partitioning on the 10X chromium controller. The Chromium Single Cell 5’ Reagent Kit with Feature Barcoding (v2) was used according to the manufacturer’s protocol. Libraries were sequences on the Illumina NovaSeq 6000 platform.

### CITE sequencing data analysis

Sequencing data was processed through the Cell Ranger Single Cell Software Suite (10X Genomics) to obtain feature-barcode matrices, then analyzed using Seurat v5. Data was first filtered to exclude cells with <250 unique features, > counts 2 standard deviations above the mean, or > 10% mitochondrial genes. RNA counts were normalized and scaled with SCTransform, and antibody-derived counts were normalized with a centered log ratio transformation. Samples were demultiplexed by organ and genotype based on HTO and ADT expression, respectively. Cells marked by two organ barcodes or two genotype barcodes were discarded as doublets. Principle component analysis was performed using the top 2000 variable features. Clustering was performed using the FindNeighbors function with 16 dimensions and the FindClusters function with a resolution of 0.6. Contaminating non-T cell clusters were excluded, and remaining cells were annotated using marker genes conserved across all identity permutations. Differentially expressed genes (DEGs) between genotypes were detected using the FindAllMarkers function and considered differentially expressed if the adjusted P value <10e^-8^ and average log2(fold change) >0.25. DEGs were visualized on a volcano plot. Trajectory analysis was performed using the slingshot^51^ package, setting “Tcf7^+^ eff” as the root cluster. The condiments^52^ workflow was used to analyze trajectory paths for differences in fate selection (*fateSelectionTest* method) and progression (*progressionTest* method) between genotypes.

### Single nuclei sample preparation

WT and KO cells were separately FACS-sorted as Thy1.2^+^CD3^+^CD8^+^CD44^high^ CD45.1 (WT) or CD45.2 (*Prdm16* KO) from lungs of X31- infected mixed bone marrow chimeric animals infected at day 8 p.i. Samples were pooled from four mice, and 250,000 live cells per genotype were submitted to the Princess Margaret Genomics Centre for processing using the 10X Genomics Chromium Single-nuclei Gene Expression platform according to the manufacturer’s protocol. Nuclei (∼5,000 per sample) were isolated using the 10X Genomics Nuclei Isolation Kit and filtered to remove debris and doublets. Individual nuclei were encapsulated in Gel Beads-in-Emulsion (GEMs) containing barcoded oligonucleotides for mRNA (poly-dT) fragments.

### snRNA-seq analysis

RNA count matrices from single-nuclei RNA-seq data were analyzed using the Seurat package in R.^53^ Data from WT and KO samples were initially filtered to retain genes expressed in ≥5 cells and cells with <10% mitochondrial genes, >2,500 reads, and >1,000 detected genes. Outliers with unusually high read counts or feature numbers were excluded using 3.5 median absolute deviations from the median via the scater package. ^54^ Doublets were identified and removed using scDblFinder.^55^ Filtered WT and KO datasets were merged, normalized, and scaled prior to calculation of module scores for cell cycle, oxidative phosphorylation, and cytoplasmic translation, using gene sets obtained from the Gene Ontology database (https://geneontology.org). SCTransform-normalized data were integrated using Canonical Correlation Analysis (CCA).^56^ Cell clustering was performed with the Smart Local Moving (SLM) algorithm^57^ at a resolution of 0.4 and UMAP dimensionality reduction was applied for visualization. Cluster-specific marker genes conserved between WT and KO samples were identified and used for annotation. Pseudobulk expression profiles were computed to assess correlations between clusters.

### Data analysis and statistics

All statistical analyses, except for bulk RNA-seq and single-cell RNA-seq, were performed using GraphPad Prism (San Diego, CA), with the specific test performed indicated elsewhere. n.s. not significant, *p < 0.05, **p < 0.01, ***p < 0.001, ****p < 0.0001 were applied.

### Data and materials availability

Transcriptomics data have been deposited in Gene expression omnibus (GEO), accession numbers GSE315337, GSE315546 for 4-1BB related data and GSE317168 and GSE317169 for Prdm16-related data. GEO data will be made available to the public upon acceptance. Flow cytometry and other data will be provided upon request.

## Results

### Intrinsic 4-1BB signaling regulates the accumulation rather than the differentiation of Ag-specific CD8 T cells during IAV infection

To investigate the T cell intrinsic role of 4-1BB in CD8 T cell responses during IAV infection, we generated mixed bone marrow (BM) chimeras in which half the hematopoietic cells expressed 4-1BB and performed CITE sequencing using the 10X Genomics platform (**Fig. 1A**). The advantage of the mixed BM chimera model is that wildtype (WT) and 4-1BB-deficient cells are exposed to the same environment, eliminating indirect effects on the T cells from higher viral load and increased inflammation in 4-1BB-deficient mice compared to WT littermates.^9^ Following reconstitution, there was a ∼1.5: 1 ratio of CD45.1 *4-1bb^+/+^* and CD45.2 *4-1bb^-/-^* T cells in the blood, consistent with 4-1BB being largely dispensable on T cells in the steady state (**Fig. 1A**). Activated donor CD8 T cells (Thy1.1^-^CD3^+^CD8^+^CD44^high^), which included a mixture of *4-1bb^-/-^* and *4-1bb^+/+^* T cells, were sorted from lung and spleen at day 8 post intranasal (i.n.) IAV infection (**Fig 1A, Supplemental Fig.1A**) for CITE-seq. Oligonucleotide-barcoded antibodies, added prior to cell sorting, were used to delineate organ source and genotype of the T cells (**Supplemental Fig.1B**). After quality control, filtering, and sample demultiplexing by organ origin, 5,404 cells from lung and 5,992 cells from spleen remained for downstream analysis (**Fig. 1B**). While the *Tnfrsf9 (4-1bb)* transcript was detected in cells from both genotypes, bulk RNA sequencing showed that *4-1bb^+/+^*cells contained mostly exonic reads of *4-1bb,* while *4-1bb^-/-^*cells contained many reads mapped to intronic regions (**Supplemental Fig.1C**), reflecting that the *4-1bb^-/-^* mice were generated through genetic disruption of the ATG start codon via insertional mutation^42^ and still retained transcription of *4-1bb*.

**Figure 1.**
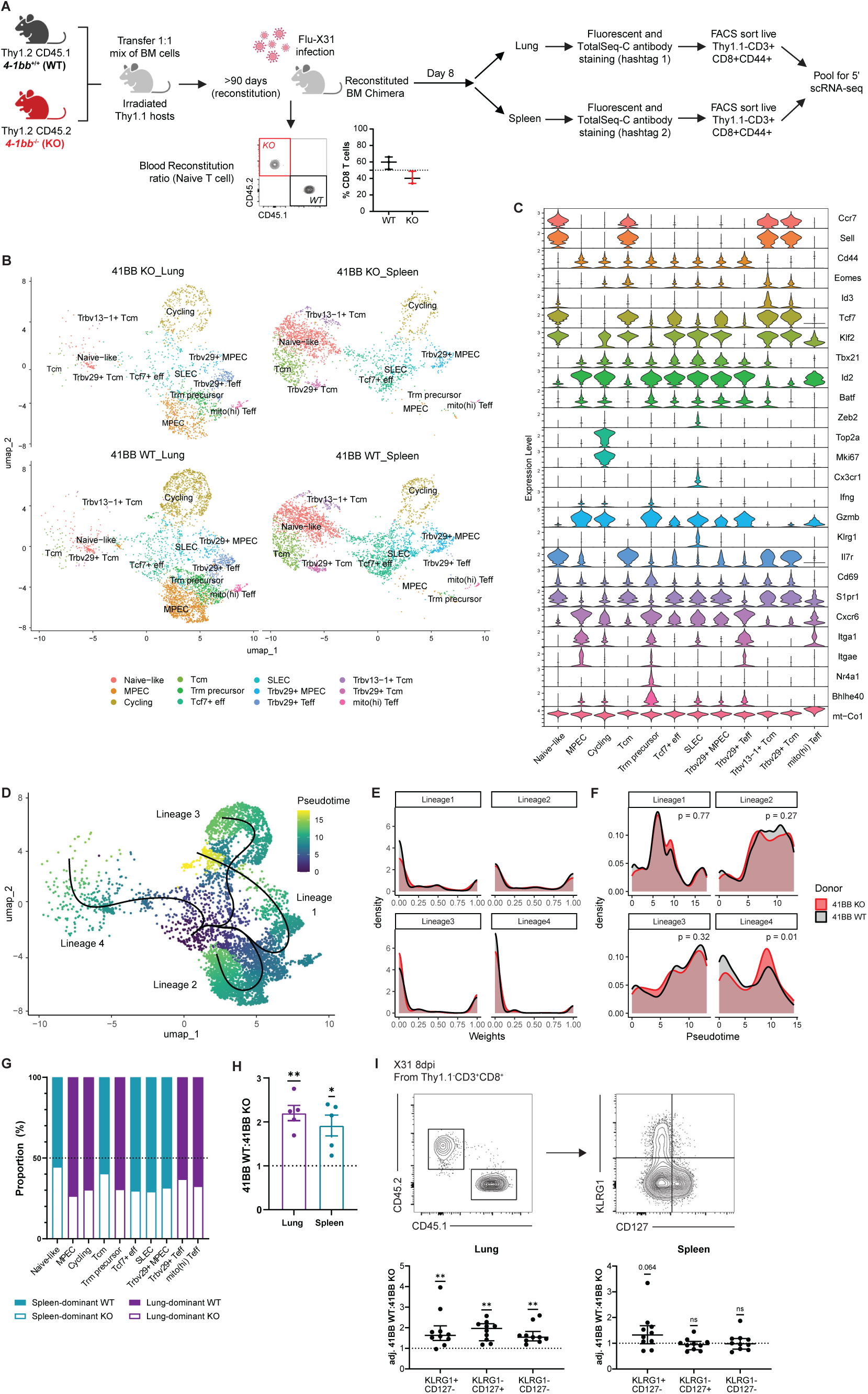
Intrinsic 4-1BB signaling controls the accumulation rather than the differentiation of CD8 T cells during IAV infection. **(A)** Experimental scheme. Lethally irradiated Thy1.1^+^CD45.2^+^ hosts were reconstituted with a 1:1 mixture of *4-1bb^+/+^* (WT;Thy1.2^+^CD45.1^+^) and 4*-1bb^-/-^* (KO; Thy1.2^+^CD45.2^+^) bone marrow cells. Chimeric mice were rested for at least 90 days and blood chimerism was assessed, followed by intranasal (i.n.) infection with influenza A X31. Single cell suspensions of spleen or lung cells from each of three mice were separately pooled and enriched for CD8^+^CD44^hi^ cells by cell sorting and processed for single cell sequencing. **(B)** UMAP of CD8 T cell populations from the lung (left) and spleen (right) of KO (top) and WT (bottom). **(C)** Expression of selected markers for cluster annotation. **(D)** Common pseudotime trajectory for all cells in the lung. (**E**) Weight and **(F)** pseudotime distributions for WT (black) and KO (red) cells across all 4 lineages. **(G)** Proportion of WT and KO cells within each cluster, colored by dominant organ source. **(H)** WT/KO ratio grouped by dominant organ source. **(I)** Representative flow cytometry gating of WT and KO CD8 effector subpopulations in mixed bone marrow chimeras 8 days p.i. with influenza-X31 and summary ratios. WT/KO ratios post infection were normalized to the pre-infection WT/KO ratio in blood CD8 T cells from the same mouse. Statistical analyses: One sample t-tests (h, i). ** p<0.01; * p<0.05.

Twelve clusters were identified across spleen and lung, based on transcriptional profiles (**Fig. 1B,C**). Naïve-like (*Cd44* low) and Tcm clusters were identified by high expression of *Ccr7*, *Sell*, *Tcf7*, and *Il7r*, and a low tissue residency profile (low *Cxcr6*, *Itga1*, *Itgae*; high *S1pr1*). A cluster of effector cells (*Gzmb*, *Id2*, *Cxcr6*) also highly expressed *Tcf7* without Tcm-related genes (*Ccr7*, *Sell*, *Il7r*), suggesting that they are stem-like but not Tcm precursors. A cluster of cycling cells was highly enriched in transcripts encoding genes related to cell proliferation (*Top2a*, *Mki67*). A small cluster of effector cells was identified with a high percentage of mitochondrial genes like *mt-Co1*. The short-lived effector cell cluster expressed *Zeb2*, *Klrg1*, and *Cx3cr1*, while memory-precursor effector cell expressed low *Klrg1* and high *Il7r*. A Trm precursor cluster was identified with high expression of the Trm transcriptional regulators *Nr4a1* and *Bhlhe40*,^58, 59^ as well as an increased tissue residency profile (high *Cd69*, *Cxcr6*, *Itga1*, *Itgae*; low *S1pr1*). Additionally, across both memory and effector clusters we observed separation of oligoclonal populations defined by expression of *Trbv29* or *Trbv13-1*, the preferential TCRβ variable chains used by D^b^/Nucleoprotein (NP) _366_-specific or D^b^/Acid polymerase (PA)_224_-specific CD8 T cells, respectively.^60, 61^

*4-1bb^+/+^ and 4-1bb^-/-^* T cells were represented in all T cell clusters within each tissue (**Fig. 1B**), suggesting that 4-1BB is dispensable for formation of all the observed T cell subsets. To further investigate whether loss of *4-1bb* influenced T cell differentiation, we performed pseudotime trajectory analysis using the slingshot^51^ and condiments^52^ workflows. A common trajectory was fitted to all cells from the lung, which identified 4 lineages emerging from the most stem-like “Tcf7^+^ effector” cluster (**Fig. 1D**). Lineages 1 and 3 represent the progression of early to activated effector cells with branches that terminate at the cycling state, whereas lineage 2 show the transition from early to late effector cells with memory potential. Lineage 4 represents the transition into Tcm, which are proportionally low in the lung during acute IAV. We did not observe any differences between *4-1bb^+/+^ and 4-1bb^-/-^* cells in fate selection (p value = 0.291, **Fig. 1E**), and minor differential progression only within lineage 4 (**Fig 1F**).

Although loss of *4-1bb* did not appear to affect the differentiation of effector T cells, *4-1bb^+/+^* cells outnumbered *4-1bb^-/-^* cells across all subsets of T cells identified (2.20-fold in lung, 1.92-fold in spleen) (**Fig. 1G,H**) despite initial reconstitution of similar numbers of *4-1bb^+/+^*and *4-1bb^-/-^* donor cells in the blood pre-infection (median ratio of 1.49:1) (**Fig. 1A**). To validate this preferential accumulation of *4-1bb^+/+^* cells across different CD8 T cell subsets, we performed flow cytometry to assess the ratio of *4-1bb^+/+^* to *4-1bb^-/-^* cells in mixed BM chimeras (**Fig. 1I**). To account for variation in reconstitution of mixed BM chimeras, data for each mouse are normalized to the pre-infection *4-1bb^+/+^:4-1bb^-/-^* ratio of CD8 T cells in the blood. Across effector subsets with a short-lived (KLRG1^+^CD127^-^), memory precursor (KLRG1^-^CD127^+^), or double-negative phenotype, by day 8 post-infection (p.i.), *4-1bb*^+/+^ cells have a significant accumulation advantage over their 4-1BB^-/-^ counterparts in the lung, confirming the differences seen based on scRNA sequencing (**Fig. 1I**). Conversely, in the spleen there was no significant difference in accumulation of *4-1bb^+/+^* over *4-1bb^-/-^* T cells, although there was a trend towards preferential accumulation of *4-1bb^+/+^* cells in the KLRG1^+^CD127^-^ subset (**Fig. 1I**). Collectively, these data suggest that 4-1BB is not a major driver of CD8 T cell differentiation during acute IAV infection but confers an accumulation advantage to the activated CD8 T cells, particularly at the site of infection.

### 4-1BB signaling during IAV infection induces Prdm16 expression in T cells

Despite the difference in accumulation of *4-1bb^+/+^* and *4-1bb^-/-^* T cells following IAV infection, we observed few transcriptional changes between *4-1bb*^+/+^ and *4-1bb^-/-^* cells in the single cell RNA-sequencing data (**Supplemental Fig.1D**). This might reflect the transient nature of these signals or the relatively low abundance or duration of endogenous 4-1BB ligand (4-1BBL) expression in the acute infection setting. Previous work identified inflammatory monocyte-derived cells as a major source of 4-1BBL during viral infection.^62^ In the lung, these inflammatory monocyte-derived cells accumulate maximally at around day 5 of IAV infection and disappear with resolution of the infection.^63^ Therefore, to capture transcriptional changes that may have occurred at an earlier time point, we conducted bulk RNA sequencing on lung CD44^high^ CD8 T cells isolated from mixed BM chimeras at day 5 post IAV infection, with the goal of detecting transcripts induced at the time of maximal 4-1BBL expression (**Fig. 2A, Supplemental Fig.1E**). A metric multidimensional scaling plot (MDS) was generated with edgeR^47^ to visualize the similarity among samples, where the distance reflects the leading log2-fold changes between samples (**Fig. 2B**). Samples tended to cluster together by genotype and separate along dimension 2.

**Figure 2.**
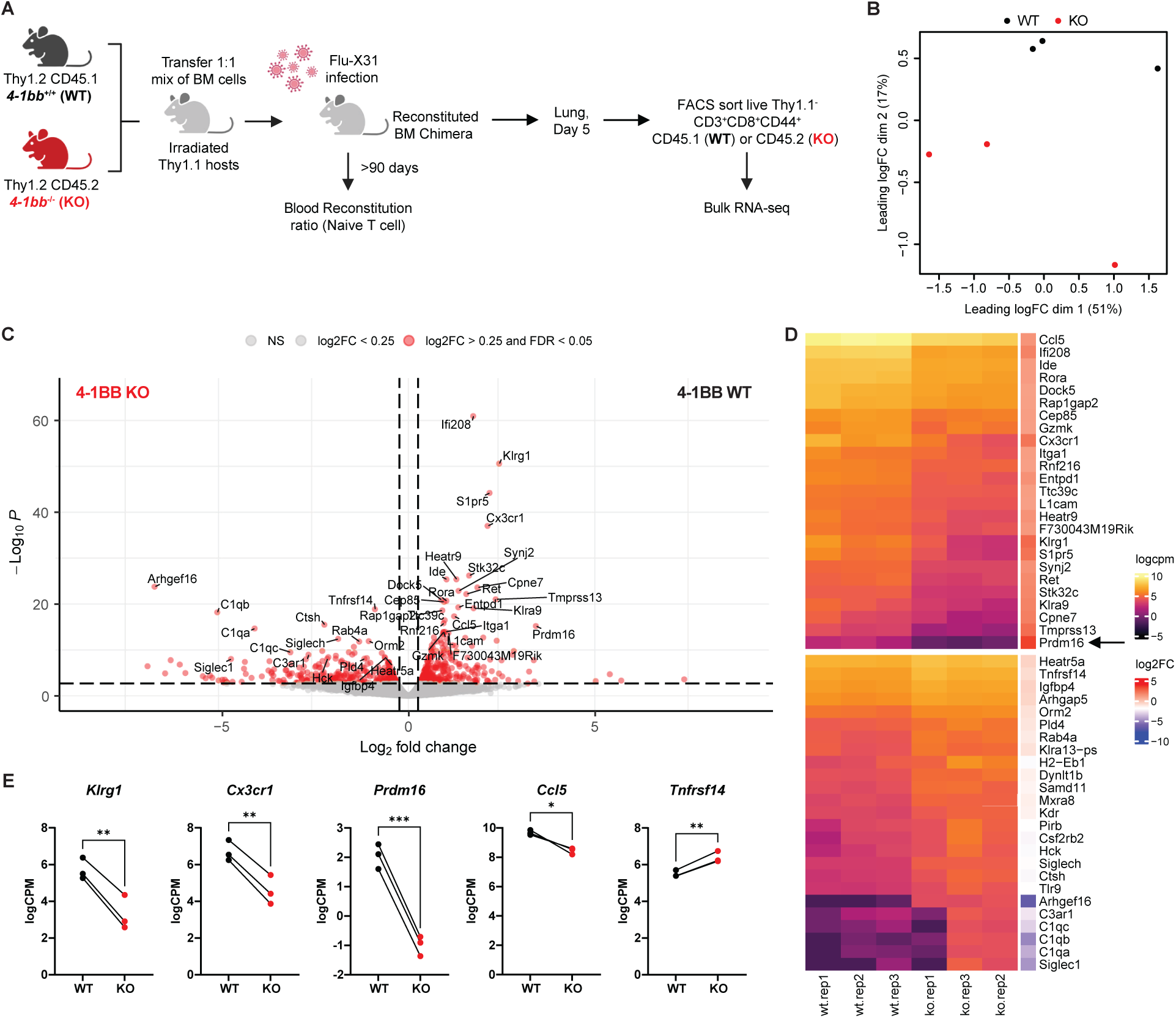
4-1BB signaling in T cells during IAV infection induces *Prdm16* expression. **(A)** Experimental schematic. Congenically distinct *4-1bb^+/+^*(WT) and *4-1bb^-/-^* (KO) CD8^+^CD44^hi^ T cells from mixed bone marrow chimeras were sorted from the lung 5 days p.i. with influenza X31 for bulk RNA sequencing with 4 mice pooled per set of WT and KO samples. **(B)** Metric multidimensional scaling plot of sequenced samples. **(C)** Differentially expressed genes (DEGs) between WT and KO samples. |log2FC| > 0.25, FDR < 0.05. **(D)** Heatmap of top 25 most significantly up-regulated and down-regulated DEGs. **e,** Log counts per million (cpm) of selected DEGs for paired samples, each derived from WT and KO cells from the same pool of chimeric mice.

Comparison of *4-1bb^+/+^* and *4-1bb^-/-^*T cells isolated from the same chimeric mice, revealed 672 differentially regulated genes (log2FC > 0.25, FDR < 0.05) (**Fig. 2C,D**). Among the most significantly upregulated genes in WT cells were those related to T cell activation (*Gzmk*, *Cx3cr1*, *Klrg1*) and migration *(Ccl5*, *Itga1*, *S1pr5*) (**Fig. 2C-E**). We also observed upregulation of *Tnfrsf14* encoding HVEM, the ligand for the inhibitory receptor BTLA,^40^ in *4-1bb^-/^*^-^ T cells (**Fig. 2C-E**), confirming the upregulation of *Tnfrsf14* in *4-1bb^-/-^* T cells that was observed in the single cell data set (**Supplemental Fig.1D**). Of the top 50 most significantly differentially regulated genes, *Prdm16* had the greatest enrichment by log2-fold change in WT over KO cells (**Fig. 2D,E**). As discussed above, Prdm16 is a transcriptional co-regulator implicated in energy metabolism in several cell types.^23, 24, 26–29, 31^ As T cell expansion and memory formation requires major metabolic reprogramming,^64^ we selected Prdm16 for further investigation.

### Prdm16 deficiency in T cells decreases influenza NP- and PA-specific CD8 T cell accumulation during IAV infection

To investigate the potential role of *Prdm16* in CD8 T cell biology, we crossed *Prdm16^fl/fl^* mice^27^ with *dLck-Cre* transgenic mice^43^ to delete *Prdm16* only in mature T cells (referred to as *Prdm16* KO throughout) (**Supplemental Fig. 2A,B**). Analysis of thymus, spleen, and inguinal lymph nodes suggested that *Prdm16* deficiency in mature T cells did not perturb thymocyte or peripheral T cell numbers in the steady state (**Supplemental Fig. 2C-E**). To examine whether *Prdm16* deficiency affects initial T cell activation, we crossed *Prdm16^fl/fl^ Lck-Cre* mice with OT-I TCR transgenic mice to generate WT and *Prdm16*-deficient OT-I T cells. Ag-stimulated OT-I WT and OT-I *Prdm16* knockout (KO) cells showed similar induction of the activation markers 4-1BB, GITR, CD69 or CD25, suggesting that *Prdm16* is dispensable for early T cell activation (**Supplemental Fig. 2F,G**).

As 4-1BB was previously implicated in CD8 T cell memory,^8, 10, 65, 66^ we infected *Prdm16* WT and KO littermates with IAV i.n. and analyzed NP- and PA-specific T cell numbers at day 30 p.i. (**Fig. 3A**). The NP-and PA-specific CD8^+^ T cell populations in the lung parenchyma included CD103^+^ and CD103^-^ Trm, as well as effector memory T cells (Tem) (**Fig. 3B,C**). At day 30 p.i., *Prdm16* KO mice displayed significantly reduced numbers of NP-specific CD103^+^ Trm and CD103^-^ PA-specific CD8 T cells with similar, albeit non-significant trends in the other lung parenchymal subsets (**Fig. 3D**). In contrast, at day 10 p.i., only the NP-specific but not the PA-specific T cells were impacted by *Prdm16* deficiency (**Supplemental Fig. 3A,B**).

**Figure 3.**
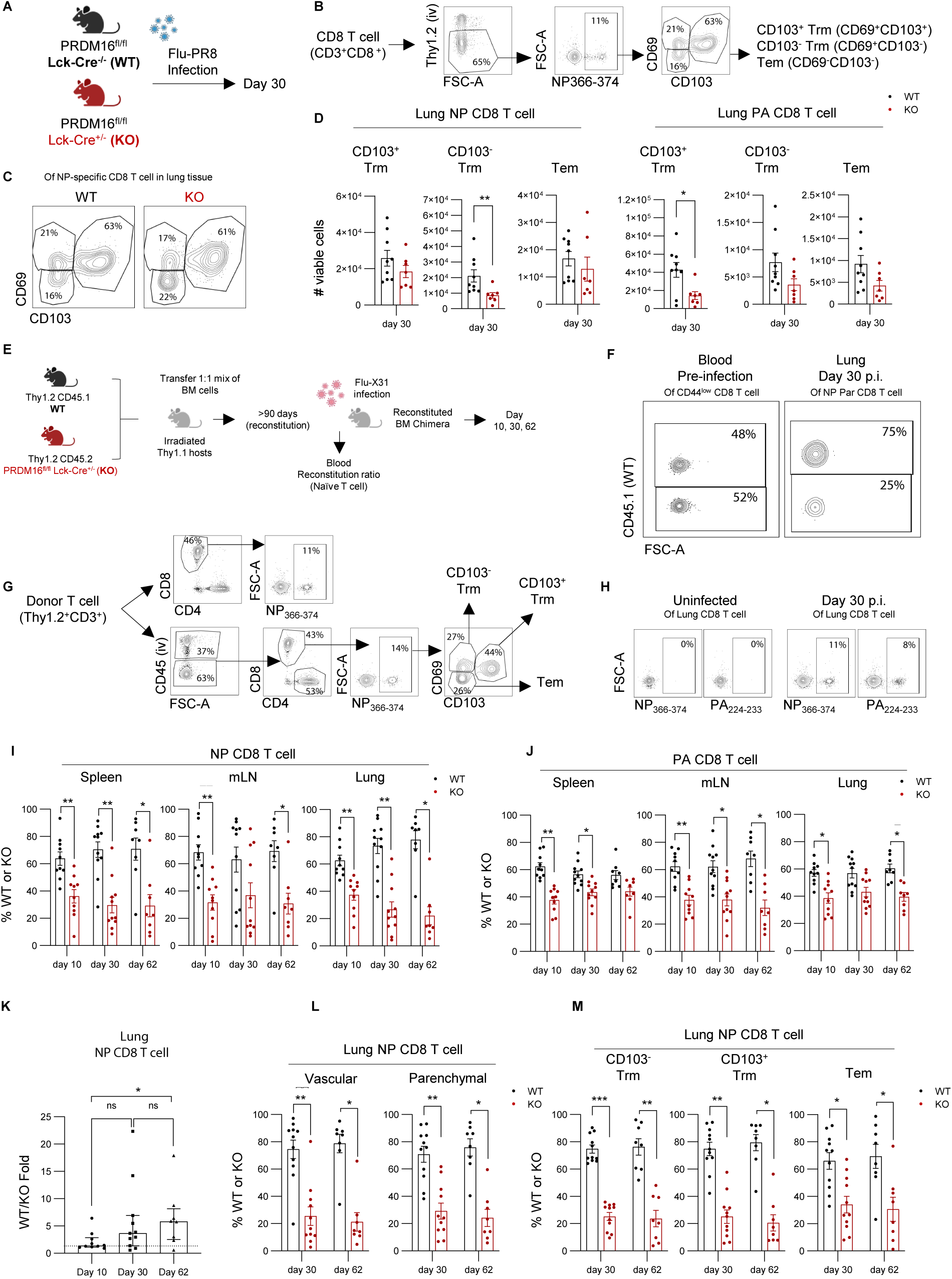
*Prdm16* deficiency in T cells decreases influenza NP- and PA-specific CD8 T cell accumulation during IAV infection. **(A)** Experimental scheme*. Prdm16^fl/fl^ dLck-Cre^-/-^* (WT) and *dLck-Cre^+/-^*(KO) littermates were infected intranasally (i.n.) with influenza A/PR8 and analyzed at 30 days post infection (p.i.). Mice were infused i.v. with FITC-anti-CD8 antibody 3 minutes prior to euthanasia, to distinguish vascular from tissue T cells.^46^ **(B)** Gating strategy for influenza nucleoprotein (NP_366–374_; NP)- or acidic polymerase protein (PA_224–233_; PA)-specific CD69⁺CD103⁺ tissue-resident memory (Trm), CD69⁺CD103⁻ Trm, and CD69⁻CD103⁻ effector memory (Tem) cells in lung parenchyma, with representative staining shown for NP-specific T cells. **(C)** Representative flow cytometry plots of WT and KO Trm and Tem subsets. **(D)** Numbers of viable NP- or PA-specific Trm and Tem populations. Data are pooled from two independent experiments (7-9 mice total). Mann-Whitney U test; **p < 0.01, *p < 0.05. Mean ± s.e.m. **(E)** Schematic of mixed bone marrow chimeras. Lethally irradiated Thy1.1^+^CD45.2^+^ hosts were reconstituted with a 1:1 mixture of bone marrow cells from WT (*Thy1.2^+^CD45.1^+^*) and KO (*Thy1.2^+^CD45.2^+^ Prdm16^fl/fl^ dLckCre^+/-^*) mice. Chimeric mice were rested for >90 days, and chimerism was assessed in blood CD8 T cells before infecting intranasally with influenza A X31. Mice were euthanized for analysis at day 10, 30, and 62 p.i. **(F)** Representative plots of WT and KO CD8 T cells in blood (pre-infection, left) and in the NP-specific lung CD8 T cell pool (day 30 p.i., right) from the same mouse. **(G-H)** Representative gating strategy (**G**) and flow plots (**H**) for NP- and PA-specific CD8 T cells in the lung. **I-J** Frequency of WT and KO cells within NP-specific (**I**) and PA-specific (**J**) CD8 T cell pools in infected lung parenchyma. **K,** WT/KO ratio in NP-specific CD8 T cells in lung parenchyma at days 10, 30, and 62 p.i. **L-M**, Frequency of WT and KO cells in NP-specific lung CD8 T cells based on vascular or parenchymal localization based on i.v. antibody labelling of vascular cells (**L**), and within Trm and Tem subsets in the lung parenchyma (**M**). WT or KO frequencies post infection were normalized to the pre-infection WT/KO ratio in blood CD8 T cells from the same mouse. Statistical analyses: Wilcoxon test (**I,J,L,M)**; pre- vs post infection) and Mann–Whitney U test (**K**; normalized WT/KO ratio across time points). Data are pooled from 8 to 11 individual chimeric mice in 2 to 3 independent experiments. Mean ± s.e.m. (**I,J,L,M**) and median ± interquartile range (IQR) (**K**). ***p < 0.001, **p < 0.01, *p < 0.05.

To compare the WT and *Prdm16^-/-^* T cells in the same inflammatory context, we generated mixed BM chimeras with an approximately 1:1 ratio of WT (CD45.1) and *Prdm16* KO (CD45.2) donor cells (**Fig. 3E-H**). Following IAV infection, WT CD8 T cells consistently outnumbered their *Prdm16* KO counterparts for both NP- and PA-specific pools, with more substantial effects observed for NP-specific CD8 T cells (**Fig. 3I,J**). This difference was observed as early as day 10 p.i., and persisted to day 62 p.i. in the spleen, mLN, and the lung. For NP-specific T cells in the lung, the effect of *Prdm16* deficiency significantly increased between d10 and day 62 (**Fig. 3K**). WT NP-specific T cells outnumbered *Prdm16* KO cells across vascular and parenchymal compartments, and across all 3 memory T cell subsets analyzed (Tem and CD103^+^ and CD103^-^ Trm) (**Fig. 3L,M),** with smaller effects on PA-specific T cells, as well as on the total CD44^high^ CD4 CD69^+^CD11a^high^ Trm population **(Supplemental Fig. 3C-E).** *Prdm16* deletion also decreased the proportion of IFN-γ- and CD107-expressing CD8 T cells following restimulation, but not the level of IFNγ or CD107 per cell (**Supplemental Fig. 3F**). Thus, Prdm16 contributes to increased numbers of Ag-specific CD8 effector and memory T cells following IAV infection, with greatest effects at the memory time point, whereas the effector function per T cell is not impacted.

### Prdm16 deficiency impairs the accumulation of TCR transgenic OT-I CD8 T cells

Next, to facilitate more in depth analysis and to assess the CD8 T cell-intrinsic role of Prdm16 under conditions of a fixed TCR, we injected congenically marked WT and *Prdm16* KO OT-I T cells at a 1:1 ratio into CD45.1/2 recipients, followed by i.n infection with IAV carrying the SIINFEKL epitope (PR8-OVA) (**Fig. 4A**). The ratio of OT-I WT to *Prdm16* KO donor cells was evaluated in the lung, mediastinal lymph node (mLN), and spleen at days 5, 8, 15, and 30 p.i (**Fig. 4B-J**). WT OT-I cells consistently outnumbered their *Prdm16* KO counterparts in the lung, with the effect significantly increasing between the effector (d5-8) and memory time points (day 30) (**Fig. 4E,F**). The effects of *Prdm16* deletion on T cell accumulation were also more pronounced in the lung compared to the draining mediastinal LN (mLN) or spleen (**Fig. 4G-I**). Within the mLN, the loss of *Prdm16* affected the Tem, but not the Tcm OT-I T cells (**Fig. 4J**) and within the lung, the effects of *Prdm16* deletion were greater on parenchymal over vascular OT-I T cells (**Fig. 4K, L**) and showed greater effects on Trm over Tem (**Fig. 4M,N**). We also observed similar enrichment of WT over *Prdm16* KO cells when Ag-specific T cells were enumerated based on frequency of degranulating (CD107a^+^) or cytokine (IFN-γ, IL-2, TNF) producing cells following restimulation (**Supplemental Fig. 4A,B**). Thus, *Prdm16* deletion decreases the frequency of responding OT-I T cells but not the levels of effector molecules per cell.

**Figure 4.**
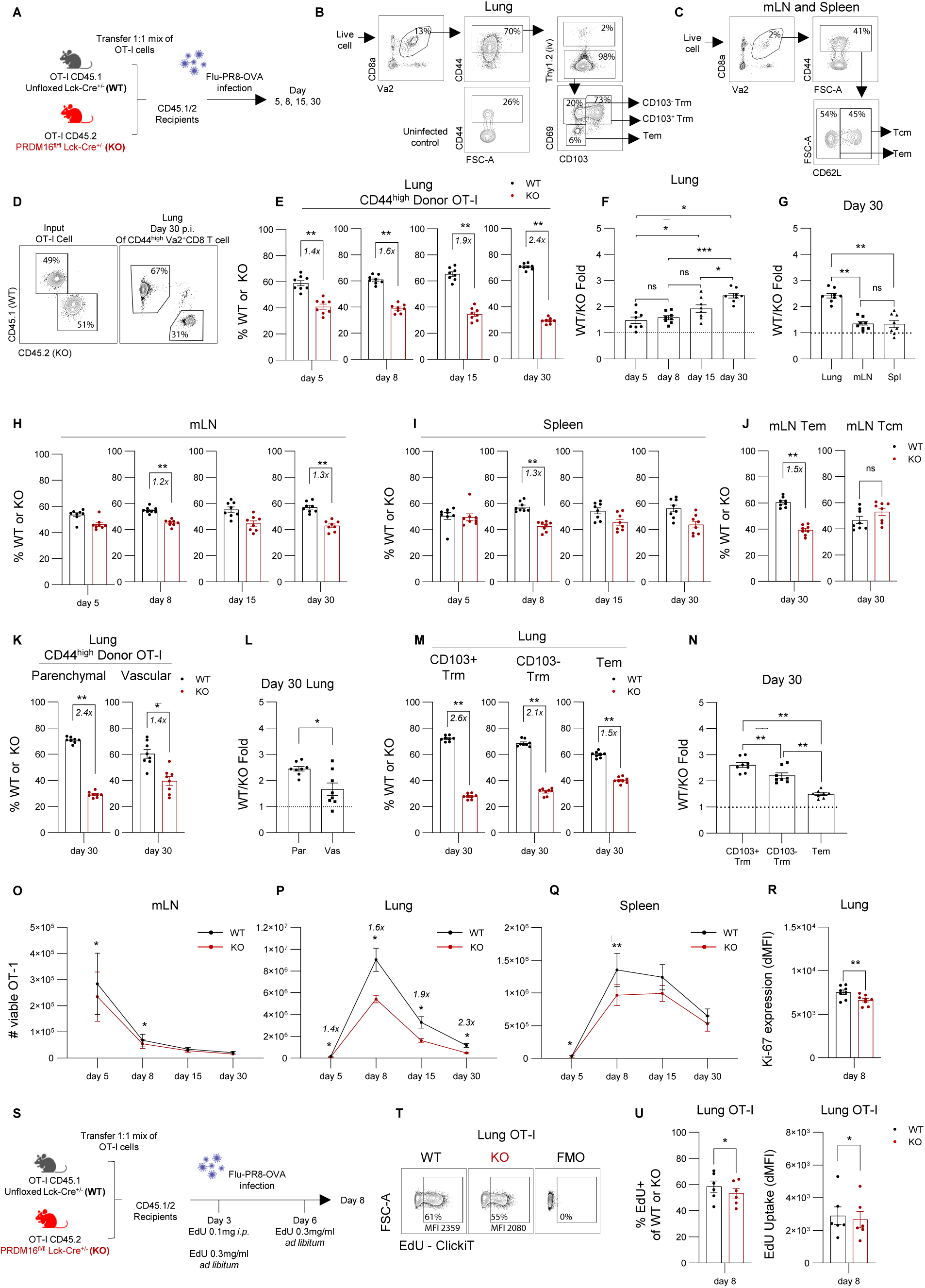
*Prdm16* deficiency decreases OT-I TCR transgenic T cell accumulation following IAV infection. **(A)** Experimental scheme. OT-I cells from spleens of OT-I CD45.1 *Prdm16^wt/wt^ dLck-Cre^+/-^* (WT) or OT-I CD45.2 *Prdm16^fl/fl^ dLck-Cre^+/-^* (KO) mice were adoptively transferred into CD45.1/2 recipients, followed by infection with Influenza A/PR8-OVA and analyzed on days 5, 8, 15, and 30 post infection (p.i.). **(B-C)** Representative gating strategy for donor OT-I cells in lung Trm and Tem subsets (**B**) or central memory (Tcm) and Tem subsets in mLN and spleen (**C**). **(D)** Representative flow cytometry plots of input WT and KO OT-I cells before adoptive transfer (left) and in lung at day 30 p.i. (right). **(E)** Frequency of WT and KO OT-I cells in the CD44^high^ donor OT-I pool in infected lung. **(F-G)** WT/KO ratio across time points in the lung (**F)** or across organs at day 30 p.i. (**G**). **(H-J)** Frequency of WT and KO OT-I cells in CD44^high^ donor pool in mLN (**H**) and spleen (**I**), and within Tem (left) or Tcm (right) subsets at day 30 p.i. (**J**). **(K)** Frequency of WT and KO OT-I cells in lung parenchymal or vascular compartment and corresponding WT/KO ratio **(L). (M)** Frequency of WT and KO OT-I cells in lung Trm and Tem subsets and (**N)** corresponding WT/KO ratio. **(O-Q),** Number of viable WT and OT-I cells in the mLN (**O**), lung (**P**), and spleen (**Q**). **(R)** Expression of Ki67 in WT and KO OT-I cells in lung at Day 8 p.i. **(S)** Experimental schematic of adoptive transfer as in **(A)**, with recipients receiving 100 μl of 1mg/ml EdU i.p. on day 3 p.i. and 0.3 mg/mL EdU in drinking water from day 3 to day 8 p.i., followed by analysis on day 8. **(T)** Representative flow cytometry plot showing EdU uptake in WT and KO OT-I cells. (**U**) Percentage of EdU⁺ cells (left) and dMFI (right) in WT and KO OT-I cells. Data are pooled from two independent experiments (6 to 8 mice) Wilcoxon test; mean ± s.e.m. ***p < 0.001, **p < 0.01, *p < 0.05.

When converted to total numbers of T cells, we observed the most substantial effects of *Prdm16* deletion on lung T cell numbers, with effects increasing slightly from the effector to memory time points (**Fig. 4O-Q**). There was a small but significant increase in the proportion of Ki67^+^ (proliferating) T cells at day 8 in WT versus KO OT-I (**Fig. 4R**). Similarly, based on EdU uptake we also observed that WT cells showed a small but significant increase in post-priming proliferation between day 3 and 8 compared to *Prdm16* KO OT-I T cells (**Fig. 4S-U**).

Taken together, the OT-I TCR transgenic T cell system corroborates observations from the endogenous response with respect to *Prdm16* expression resulting in increased effector and memory T cell numbers without impacting cytokine production per cell. The results also show that deletion of *Prdm16* has greater effects in the lung parenchyma than the lung vasculature or secondary lymphoid organs, with a small but significant decrease in proliferation of *Prdm16 KO* OT-I T cells.

### Prdm16 overexpression increases OT-I TCR transgenic T cell accumulation following IAV infection

We next investigated the effect of *Prdm16* overexpression on the OT-I T cell response. *In vitro* activated OT-I cells were transduced with retroviral vectors encoding Prdm16 and EGFP (OE) or EGFP alone (mock), expanded in cytokines, sorted based on GFP expression and then transferred in equal numbers into CD45.1/2 recipients, followed by i.n. PR8-OVA infection (**Fig. 5A, B**). OE cells consistently displayed a competitive numerical advantage over mock-transduced cells as early as day 8 in the lung and this difference persisted to day 30, with similar fold differences observed in the mLN and spleen at day 8, 15, and 30 (**Fig. 5C-F**). OE OT-I outnumbered mock-transduced OT-I cells in both parenchymal and vascular compartments and across effector and memory T cell subsets in lung, spleen and mLN (**Fig. 5G-I**). Similar effects were observed based on analyzing degranulation (CD107) or cytokine production (IFNγ, IL-2) following restimulation at day 30 (**Supplemental Fig. 4C, D**). These results indicate that enforced *Prdm16* expression provides a competitive numerical advantage to effector and memory T cells, following IAV infection *in vivo*, independent of T cell subset or location analyzed, without affecting the level of effector molecules per cell.

**Figure 5.**
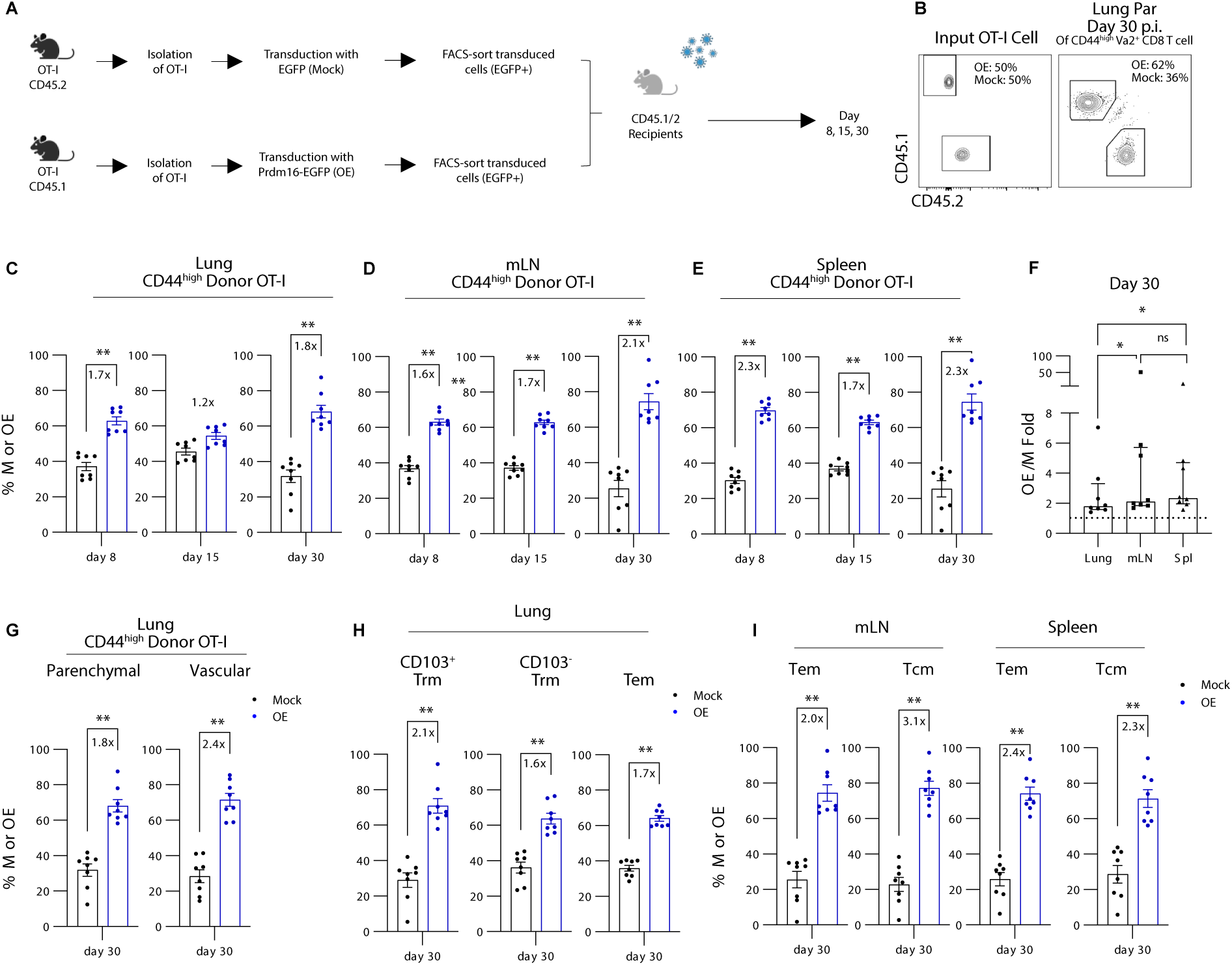
*Prdm16* overexpression increases OT-I TCR transgenic T cell accumulation following IAV infection. **(A)** Experimental schematic. As described in detail in the methods, CD45.1 or CD45.2 OT-I cells were isolated from spleens of OT-I mice, activated with anti-CD3 and anti-CD28, treated with IL-2 for 2hrs then transduced with gamma-retrovirus vectors encoding EGFP alone (Mock, M) or *Prdm16*-EGFP (Overexpression, OE). Transduced cells were expanded in IL-2 and IL-7 for 8 days and then EGFP^+^ cells were FACS-sorted, mixed in a 1:1 ratio, and adoptively transferred into CD45.1/2 recipients, followed by influenza A/PR8-OVA infection. Cells were analyzed on days 8, 15, and 30 p.i. **(B)** Representative flow cytometry plots showing input ratio before adoptive transfer (left) and post-infection ratio in lung parenchyma at Day 30 (right). **(C-F)** Frequencies of Mock or OE OT-I cells in the lung (**C**), mLN (**D**), and spleen (**E**) over time, with corresponding OE/Mock fold ratios at day 30 (**F**). **(G)** Frequencies of Mock or OE cells in lung parenchymal versus vascular compartments at day 30 (**H**) Frequencies of Mock or OE cells in lung parenchyma, gated on Trm and Tem subsets at day 30. **(I)** Frequencies of M or OE cells in mLN or spleen, gated on Tem and Tcm subsets at day 30. Data are pooled from two independent experiments which included switched congenics (experiment 1: OT-I CD45.2 cells transduced with Mock and CD45.1 with OE; experiment 2: CD45.1 Mock and CD45.2 OE). Post-infection frequencies were normalized to the input OE/Mock ratio. Statistical analysis: Wilcoxon test comparing Mock versus OE frequencies in **C-E**, **G-I** and OE/M fold ratios in **F**. *** p<0.001; ** p<0.01; * p<0.05. Data are presented as mean ± s.e.m. (c-e, g-i) or median ± IQR (f).

### Prdm16 regulates genes involved in mitophagy and immune modulation

To investigate the transcriptional programs induced downstream of Prdm16 in T cells, we performed bulk RNA-sequencing on WT and *Prdm16* KO OT-I T cells adoptively transferred into CD45.1/2 recipients at a 1:1 ratio, and sorted from mouse lungs at day 5, 8 and 30 following IAV infection (**Fig. 6A**). As expected, *Prdm16* mRNA was higher in WT compared to KO T cells at all time points, confirming its expression in WT T cells (**Fig. 6A, B**). *Prdm16* KO OT-I T cells had reduced expression of two genes related to mitophagy, *Pink1* (encoding PTEN induced novel kinase 1) (**Fig. 6A,B**)^67^ and *Park7 (*encoding the DJ-1 protein) (**Fig. 6B**).^34, 68^ There was also a trend toward higher *Mfn2* (encoding Mitofusin 2)^69^ in *Prdm16*-expressing OT-I T cells at all 3 time points analyzed but the small increases in WT over KO did not reach significance (**Fig. 6B**). We also observed *Prdm16*-dependent expression of the small GTPase *Rab6b*,^70^ involved in vesicular trafficking from the golgi (**Fig. 6A, B**). Genes upregulated in *Prdm16* KO cells included *Btg2,* involved in T cell quiescence,^71^ *Cap1,* involved in actin regulation,^72^ and *Camk2b,* encoding calmodulin dependent protein kinase 2 **(Fig. 6A, B)**. As had been seen in *4-1bb^-/-^* T cells, *Tnfrsf14* was upregulated in *Prdm16* KO cells at all 3 time points, and its higher surface expression in *Prdm16* KO memory T cells was confirmed at the protein level (**Fig. 6A-D**). Gene set enrichment analysis (GSEA) suggests that WT cells have enrichment of hydrogen peroxide metabolic processes compared to KO cells, whereas pathways associated with negative regulation of adaptive immune responses were preferentially enriched in *Prdm16* KO cells, largely driven by HVEM (**Fig. 6E**).

**Figure 6.**
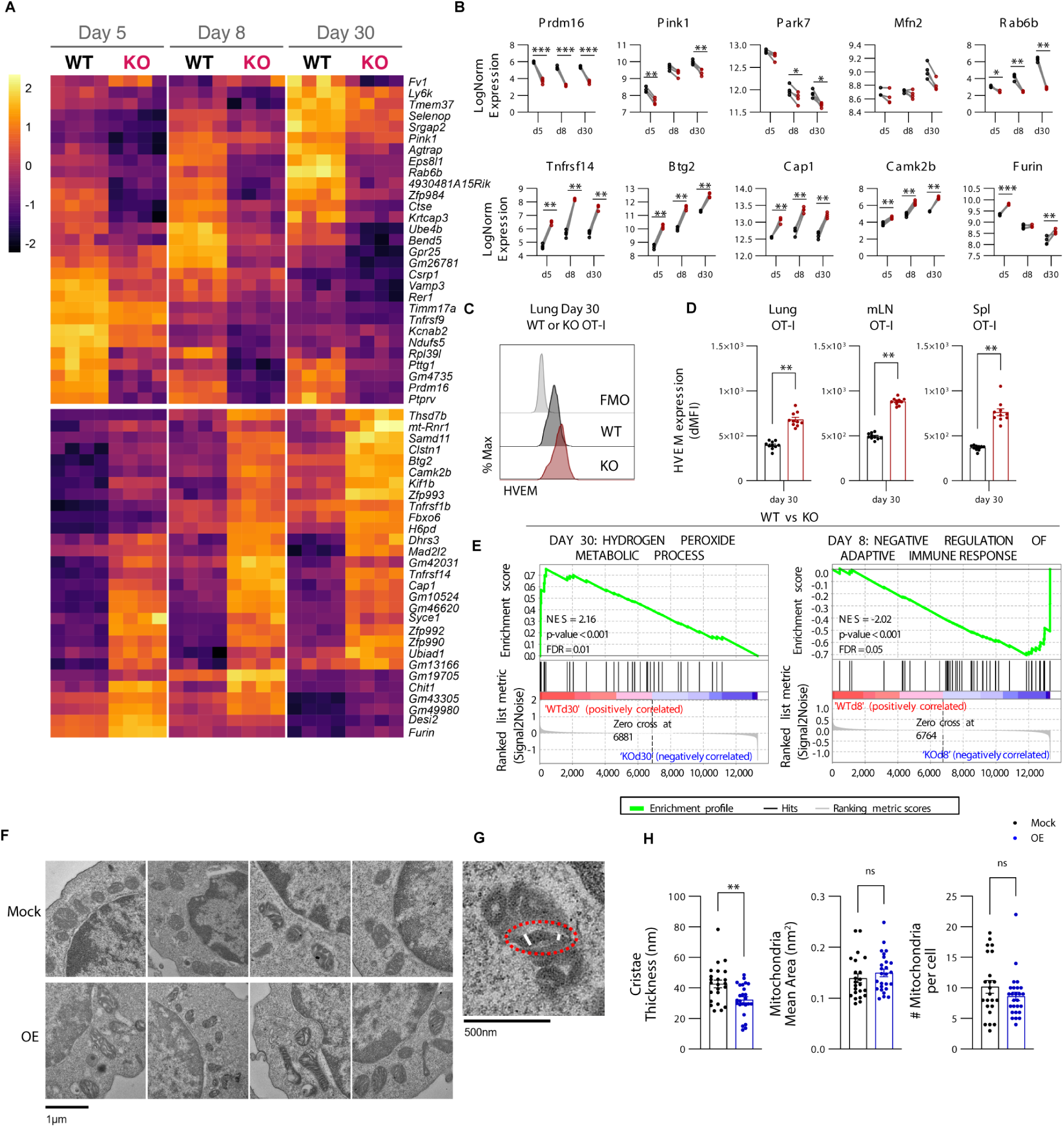
Prdm16 regulates genes involved in mitophagy and its expression is associated with more compact mitochondrial cristae. *CD45.1 Prdm16^wt/wt^ dLck-Cre^+/-^* (WT) and *CD45.2 Prdm16^fl/fl^ dLck-Cre^+/-^*(KO) OT-I cells (1:1 mix of 100,000 per genotype), were adoptively transferred in CD45.1/2 C57BL/6 recipients. One day later, mice were infected with influenza A/PR8-OVA. CD8^+^CD44^high^ CD45.1 and CD45.2 OT-I cells were sorted from lungs of 8 mice at days 5, 8, and 30 p.i. and RNA was separately isolated from each sample for bulk RNA-sequencing. **(A)** Heatmap of differentially expressed genes in WT and KO at different time points (top genes with lowest p.adj values, |log2FC| > 0.25, p.adj < 0.05), with each column representing data from one mouse. **(B)** Log normalized expression of selected genes. FDR values < 0.05 for each comparison are indicated. **(C-D)** HVEM protein expression on WT or KO OT-I cells (lung day 30 p.i.) was measured by flow cytometry with representative histograms shown in (**C**) and dMFI shown in (**D**). Data are pooled from 8 individual mouse across 2 independent experiments. Wilcoxon test; mean ± s.e.m. ***p < 0.001, **p < 0.01, *p < 0.05. **(E)** GSEA enrichment plots comparing WT and KO OT-I cells. **(F-H)** Electron microscopy (EM) analysis of Mock-transduced (Mock) or *Prdm16*-Overexpressing (OE) T cells. The Mock and *Prdm16* OE OT-I cells had been stimulated, transduced and expanded for 8 days as describe in Fig. 5a and in the methods. **(F)** Representative EM images with top 4 images from Mock transduced OT-I and lower 4 images from *Prdm16* OE cells. **(G)** Cristae measurement (2-3 cristae per mitochondrion, average of thickest and thinnest points). **(H)** Cristae thickness (left), mean mitochondrial area (middle), and mitochondrial number per cell (right); each dot represents one cell (23–26 cells from two independent experiments). Electron micrograph analyses were blinded. Mann-Whitney test; **p<0.01. Data shown as mean

### Prdm16 expression is associated with more compact mitochondrial cristae

Based on the decreased expression of *Park7* and *Pink1* in *Prdm16* KO cells, we next asked whether *Prdm16* affects mitochondrial content or ultrastructure. We compared mock-transduced and *Prdm16*-overexpressing (OE) OT-I cells by transmission electron microscopy (TEM). Although the number and area of mitochondria per cell was indistinguishable between mock and *Prdm16* OE OT-I cells, *Prdm16* OE OT-I cells exhibited more compact and organized cristae (**Fig. 6F-H**), which has been associated with more efficient electron transport.^38^ Based on flow cytometric assays, Mock and *Prdm16* OE OT-I cells showed similar mitochondrial mass and membrane potential when analyzed *in vitro* (**Supplemental Fig. 5A, B**). Similarly, flow cytometric analysis revealed no significant differences in mitochondrial mass, mitochondrial membrane potential, mitochondrial ROS or fatty uptake between WT and *Prdm16* KO OT-I T cells following IAV (PR8-OVA) infection of mice, and measured directly *ex vivo* (**Supplemental Fig. 5C-F**).

### Prdm16 drives the accumulation of effector CD8 T cells with a gene signature suggesting high ribosomal and mitochondrial activity

To examine the effect of *Prdm16* on transcriptional programs at the single cell level we performed single-nuclei sequencing (10x Genomics). WT and *Prdm16^fl/fl^ dLck-Cre*^+/-^ KO CD8 T cells were isolated from the lungs of mixed bone marrow chimeras at day 8 p.i. with IAV and WT and KO T cells sorted for analysis based on congenic markers. Gene expression analysis revealed multiple transcriptionally distinct effector CD8 T cell clusters (**Fig. 7A, B; Supplemental Fig. 6A**). We identified a distinct cluster which we termed the High_Ribo_Mt_Activity cluster, characterized by high expression of genes encoding mitochondrial and ribosomal proteins. Although clearly T cells based on expression of T cell genes such as *Cd3e* (data not shown) and *Cd8a* (**Supplemental Fig. 6A)**, this cluster did not fit the canonical profile of established T cell subsets, suggesting that it reflects a cellular state rather than a distinct lineage. Correlation analysis positioned this cluster closest to short-lived effector and proliferating effector populations (**Fig. 7C**). Consistent with its transcriptional profile, this subset exhibited high oxidative phosphorylation and protein translation scores (**Fig. 7D**) compared to other subsets and showed moderate proliferative signatures. Notably, WT cells were markedly enriched in this cluster compared to *Prdm16*-deficient cells (**Fig. 7E**). Within the cluster, WT cells displayed higher S-phase and G2/M scores, suggesting enhanced proliferative activity relative to KO counterparts (**Fig. 7F**). Other subpopulations identified include short-lived effector cells characterized by the expression of effector genes *Ccl4*, *Ifng*, and *Tnfrsf9*, as well as exhaustion markers *Havcr2*, *Lag3*, and *Tox*; trafficking effector T cells (Teff) showing higher *Klf2* expression; proliferative effector T cells expressing cell cycle markers like *Mki67*, *Pcna*, and histone genes; pre-Tcm cells expressing memory markers such as *Sell*, *Ccr7*, *Tcf7*, and *Lef1*; a pre-Trm population with classical tissue-resident cell makers such as *Itga1*, *Itgae*, and *Cxcr6*; and another pre-Trm-like population with higher expression of *Ctla4, Icos*,^73^ and ISGs (**Fig. 7G; Supplemental Fig. 6B)**. Differential gene expression analysis further identified genes that were upregulated or downregulated in patterns consistent with those observed in the bulk RNA-seq. Specifically, *Pink1* was more highly expressed in WT compared to *Prdm16* KO cells, with the most substantial effects observed in the WT High_Ribo_Mt_Activity T cells, followed by the WT pre-Trm cells. *Tnfrsf14* was most upregulated in KO pre-Trm cells (**Fig. 7G**). These results suggest that Prdm16 promotes the accumulation of T cells in a metabolically active state and that the Prdm16-induced transcriptional modifications are variable among the different CD8 T cell populations/states, with greatest effects in the High_Ribo_Mit_Activity, Pre-Trm and proliferative clusters and minimal effects in short lived effectors.

**Figure 7.**
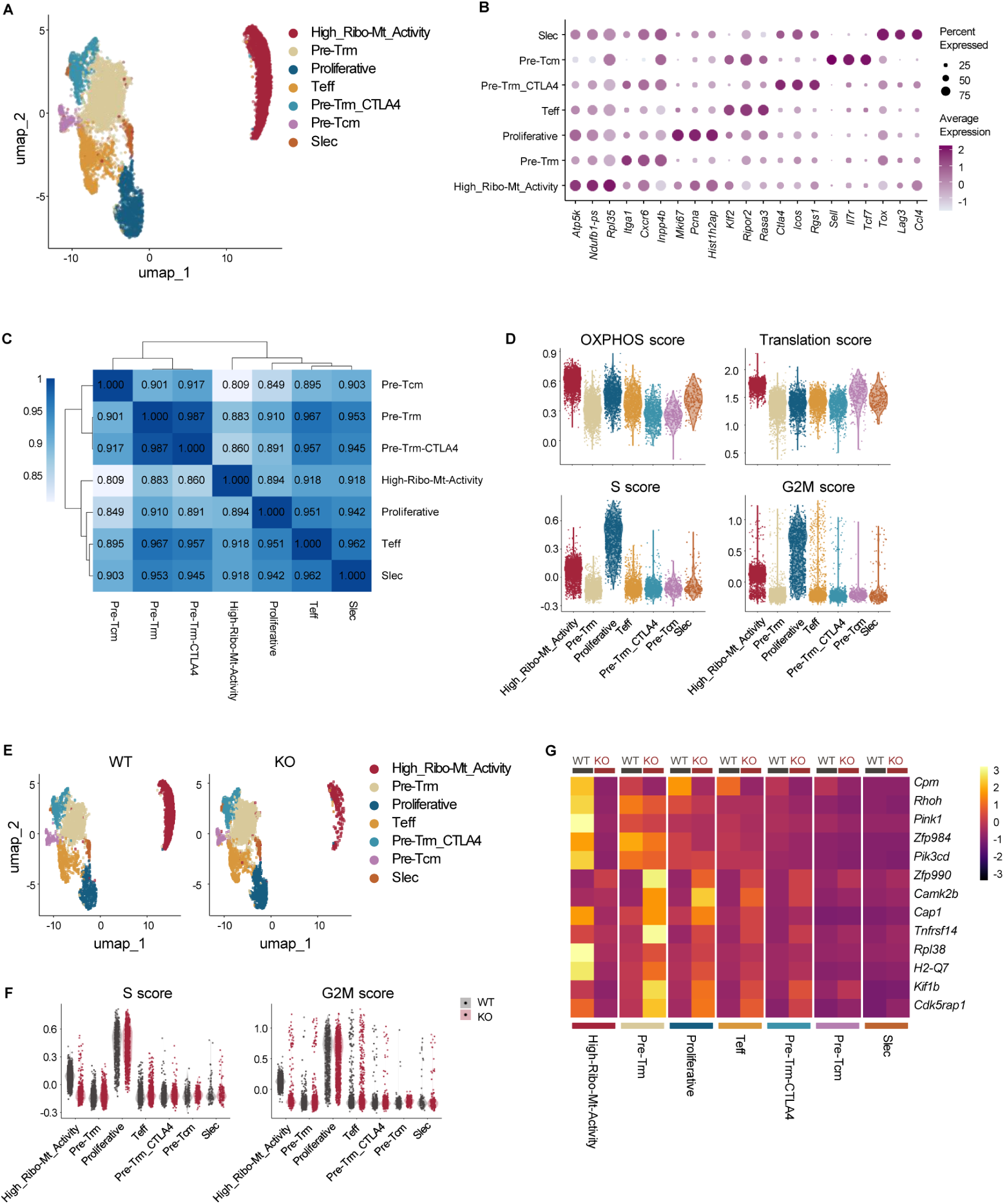
Prdm16 drives the accumulation of effector CD8 T cells with a gene signature suggesting high translational and mitochondrial activity. Lethally irradiated mice were reconstituted with CD45.1 (WT) and CD45.2 *Prdm16^fl/fl^ dLck-Cre^+/-^* (KO) bone marrow (1:1 ratio), then infected with influenza A/X31 as described above. At day 8 post infection, CD8^+^CD44^high^ CD45.1 and CD45.2 cells were separately FACS-sorted from the lungs and analyzed by snRNA-seq. **(A)** UMAP showing the clusters detected from the integrated WT and KO cells. **(B)** Dot plot showing top markers for each annotated cluster and their average expression. **(C)** Correlation heatmap for the pseudo-bulk gene expression of each cluster. **(D)** Scores for oxidative phosphorylation (OXPHOS), cytoplasmic translation, and cell cycle phases (S and G2M) of each cluster calculated from the gene expression. **(E)** UMAP plot of the clusters split by genotype origin. (**F**) Cell cycle phases score in each cluster split by genotype. **(G)** Heatmap showing the average expression of top differentially expressed genes between WT and KO for each cluster. It is important to note that our quality control pipeline excluded cells with more than 10% mitochondrial genes, eliminating most dead or dying cells from our analysis.

## Discussion

In this report we found that T cell intrinsic 4-1BB signaling drives the accumulation rather than the differentiation of CD8 T cells during IAV infection, with greater effects on T cell numbers in the lung over the spleen. The transcriptional coregulator *Prdm16* was one of the most significantly upregulated transcripts induced downstream of 4-1BB in lung T cells during IAV infection. *Prdm16* was previously noted to be upregulated in CD4 T cells by the TNFR superfamily member GITR (*Tnfrsf18*) during chronic LCMV infection,^62^ and *Prdm16* expression was also observed in CD4 effectors following LCMV Armstrong infection.^74^ However, to our knowledge the role of Prdm16 in αβ T cells has not been investigated until now. *Prdm16* deletion as well as enforced expression of *Prdm16* in T cells support a role for *Prdm16* in increasing T cell accumulation during clonal expansion and memory formation. Although both effector and memory T cell subsets were impacted by loss of *Prdm16*, *Prdm16* deficiency in OT-I T cells had larger effects on T cells in the lung parenchyma compared to the vasculature, spleen or LN and more pronounced effects on memory compared to effector T cells, with CD103^+^ Trm phenotype cells most affected. The increased effect of *Prdm16* deficiency in the lung and on resident T cell populations might reflect higher induction of *Prdm16* in the lung tissue, consistent with a greater role for TNFR superfamily effects in tissues over secondary lymphoid organs,^5^ although this was not addressed here. However, the differential magnitude of the *Prdm16* effect between secondary lymphoid organs and lung was not as apparent when looking at endogenous NP and PA-specific responses in mixed chimeras, perhaps reflecting different sensitivities of different epitope-specific T cells. Complementing the results obtained with *Prdm16* deletion, enforced expression of *Prdm16* increased CD8 T cell accumulation across tissues and subsets. Although *Prdm16* can be expressed in different isoforms,^29^ the finding that enforced expression of full length *Prdm16* shows the inverse effect of *Prdm16* knockout, suggests that the effects of *Prdm16* on T cell accumulation are largely due to the full-length form.

*Prdm16* deletion was found to have a small but significant effect on post-priming T cell proliferation from days 3-8 p.i. We focused on this time-period based on prior work defining the time course of TNFSF ligand induction on monocyte derived cells and maximal monocyte accumulation during IAV infection.^63^ Consistently, we observed a population of proliferating cells in the single nuclei RNA-seq data from day 8, referred to as the High_Ribo_Mt_Activity cluster and more strongly represented in WT compared to *Prdm16^-/-^* T cells. Of note, *Btg2,* whose gene product is involved in T cell quiescence,^71^ was upregulated in *Prdm16*^-/-^ cells, consistent with reduced proliferation. However, this small effect on proliferation may not be sufficient to explain the increases in T cell numbers over the same time span, suggesting that survival of the T cells during their expansion phase is likely also impacted.

One prominently upregulated gene in *Prdm16^-/-^* T cells was *Tnfrsf14,* encoding HVEM. Although HVEM is a TNFR superfamily member that binds the TNFSF molecule LIGHT, it also binds the inhibitory receptors BTLA and CD160.^41, 75^ On T cells, HVEM and BTLA can bind in cis to inhibit T cell responses.^39^ Knockout of HVEM in T cells results in increased T cell responses and exacerbated autoimmunity, suggesting a primary role for HVEM as a negative regulator of T cells.^40^ Of note, HVEM was also noted to be upregulated in *4-1bb*^-/-^ T cells in both single cell and bulk RNA sequencing data sets (this report) as well as in *Gitr*^-/-^ CD4 T cells during chronic LCMV infection.^62^ Thus, Prdm16 may increase Ag-specific CD8 T cell numbers in part by limiting HVEM expression downstream of TNFR superfamily members.

Transcripts of genes involved in mitochondrial quality control, including *Pink1* and *Park 7* were detected at higher levels in WT compared to *Prdm16*^-/-^ T cells at multiple time points in the bulk RNA-seq data. Similarly, in the single nuclei data, *Pink1* was also higher in WT compared to KO cells, with highest expression in the High_Ribo_Mt_Activity WT cluster, followed by the pre-Trm WT cluster. *Park7* (encoding DJ-1) was detected at higher levels in WT compared to *Prdm16^-/-^* cells at all 3 time points in the bulk RNA_seq data, but with decreasing expression from effector to memory time points. Loss of function mutations in *Pink1* and *Park7* have been well studied for their role in autosomal recessive Parkinsons disease and both Pink1 and DJ-1 (encoded by *Park7)* have been implicated in the Pink1/Parkin/DJ-1 mitophagy pathway.^35, 68, 76^ Recent evidence shows mitophagy is upregulated in T cells during anti-viral responses and that Parkin deficiency impairs memory T cell formation.^36^ Consistently, Parkinsons disease patients showed perturbations in CD8 T cell subsets and defects in Ag-specific CD8 T cell responses.^36^ In our study, *Pink1* was expressed more highly in memory T cells compared to the early effector time point based on bulk RNA-seq. Therefore, we hypothesize that Prdm16 increases effector and memory T cell numbers in part through effects on induction of mitophagy pathway proteins to maintain mitochondrial quality during T cell expansion and again during the memory phase of the response. We investigated mitochondrial quality by several approaches. We found that OT-I cells with or without *Prdm16* overexpression had similar numbers and area of mitochondria based on TEM. However, mitochondria from OT-I cells overexpressing *Prdm16* had more compact cristae compared to mock transduced OT-I, which has been associated with more efficient electron transport.^37, 38^ It is plausible that by removing the damaged mitochondria, mitophagy enriches for mitochondria with more compact and well-formed cristae.

Flow cytometry-based assays showed similar mitochondrial mass, mitochondrial ROS, or mitochondrial membrane potential with or without *Prdm16*. Several studies have shown that supraphysiological 4-1BB signaling using agonistic antibodies or CAR-T containing 4-1BB signaling domains increases mitochondrial fusion and biogenesis,^17–19^ with one study showing that these increases involve a signaling axis including p38 mitogen activated protein kinase, activating transcription factor 2 (ATF2) and Peroxisome proliferator-activated receptor-gamma coactivator (PGC)1α.^18^ Interestingly, in brown fat, Prdm16 regulates PGC1α transcription,^77^ thus it is possible that these two pathways downstream of 4-1BB could interact to optimize mitochondrial function. In HSC, Prdm16 induces *Mfn2,* to induce mitochondrial fusion.^28^ Although we saw small increases in *Mfn2* at all 3 time points in WT compared to *Prdm16*^-/-^ T cells in the bulk RNA-seq data, these differences were very small and did not reach significance.

A limitation of our study is the subtle effect of Prdm16 at single gene levels, limiting the number of DEGs detected. However, the sum of these effects can clearly double the number of Ag-specific effector and memory T cells over the course of the T cell response to IAV.

In conclusion, the data presented here showed that during respiratory IAV infection, 4-1BB in T cells induces *Prdm16*, which in turn contributes to increasing the number of antigen-specific T cells with effects persisting from the effector to the memory phase of the response. Expression of *Prdm16* is associated with decreased expression of the inhibitory ligand HVEM, and the induction of *Pink1* in cells with high mitochondrial and protein translation activity as well as in pre-Trm. *Prdm16* overexpression was associated with more compact mitochondrial cristae, suggesting a role for Prdm16 in mitochondrial quality control in T cells. Taken together, our work identifies a 4-1BB-Prdm16 axis that is induced in T cells during viral infection to support T cell accumulation and memory formation.

## CRediT authorship contribution statement

Conceptualization: SL, KM, CdAH, THW ; Obtained funding: THW; Wrote and edited manuscript: SL, KY, CdAH, RE, THW; Conducted experiments and analyzed data: SL, KY, CdAH, RE and AD.

## Acknowledgments

We thank Birinder Ghumman for technical assistance, Nathalia Viera Batista for help with initial Prdm16 mouse crosses; Natalie Simard for advice on flow cytometry; Troy Ketela and the Princess Margaret Genomics facility for transcriptomics analysis, the National Institute of Health (NIH) Tetramer Core Facility at Emory University for MHC tetramers, and Michele Anderson, Matthew Buechler, Thierry Mallevaey for critical review of the manuscript. This work was funded by Canadian Institutes of Health Research Grants FDN-143250 and PJT-178020 to THW. THW holds the Canada Research Chair in Anti-viral immunity at the University of Toronto. KY was funded in part by an Ontario Graduate scholarship; KY and SY were funded in part by an Emerging and pandemic infections consortium doctoral awards at the University of Toronto.

## Appendix A Supplementary data

## Abbreviations

Ag: antigen
BM: bone marrow
BTLA: B and T lymphocyte attenuator
CAR: chimeric antigen receptor
HSC: hematopoietic stem cell
HVEM: Herpes virus entry mediator
IAV: influenza A virus
i.n.: intranasal
KO: knockout
mLN: mediastinal lymph node
NP: nucleoprotein
OE: overexpression
PA: acidic polymerase
p.i.: post infection
Prdm16: PR domain containing 16
PR8: Influenza A/PR8 H1N1 1934
TCID_50_: tissue culture infectious dose 50
Tem: effector memory T cells
Tcm: central memory T cells
Trm: resident memory T cells
WT: wildtype

