## Supplemental material for "T cell intrinsic 4-1BB signals induce Prdm16 to increase effector and memory T cell numbers during respiratory influenza infection"

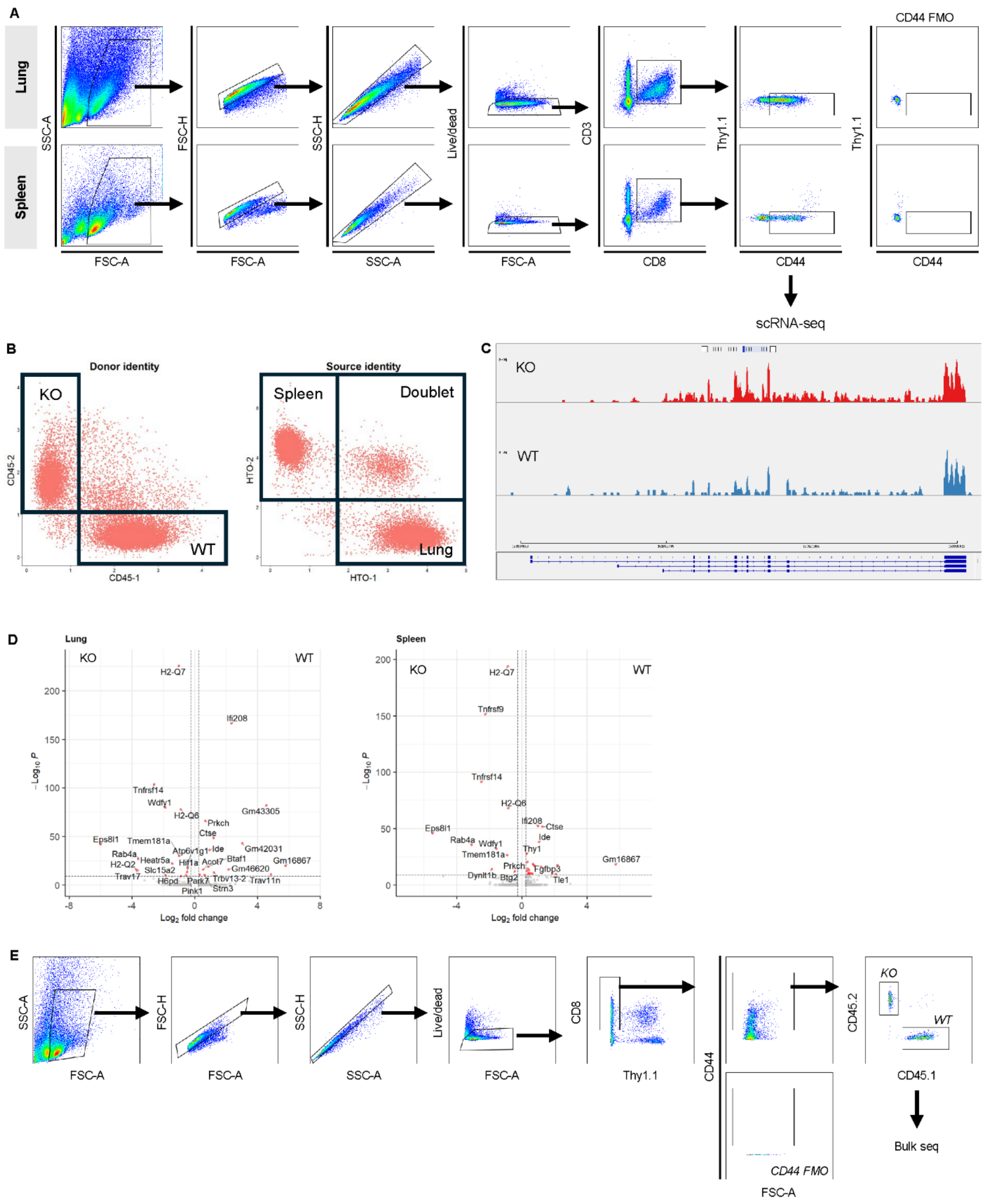


**Supplemental Figure 1. Transcriptomics analysis of 4-1BB$^{+/+}$ and 4-1BB$^{-/-}$ T cells.** **(A)** Representative flow cytometry gating for sorting of samples for single-cell sequencing. **(B)** Sample deconvolution from hashtag oligos (HTO-1, HTO-2) and antibody-derived tags (CD45.1, CD45.2) for 4-1BB single cell RNA-sequencing experiment. **(C)** Sashimi plot showing read coverage over the *Tnfrsf9* (*4-1bb* gene for WT (blue) and KO (red) samples from bulk RNA sequencing. **(D)** Volcano plot of differentially expressed genes in lung and spleen based on analysis of all WT and KO cells in the single cell RNA-seq data. **(E)** Representative flow cytometry gating for sorting of samples for bulk sequencing.

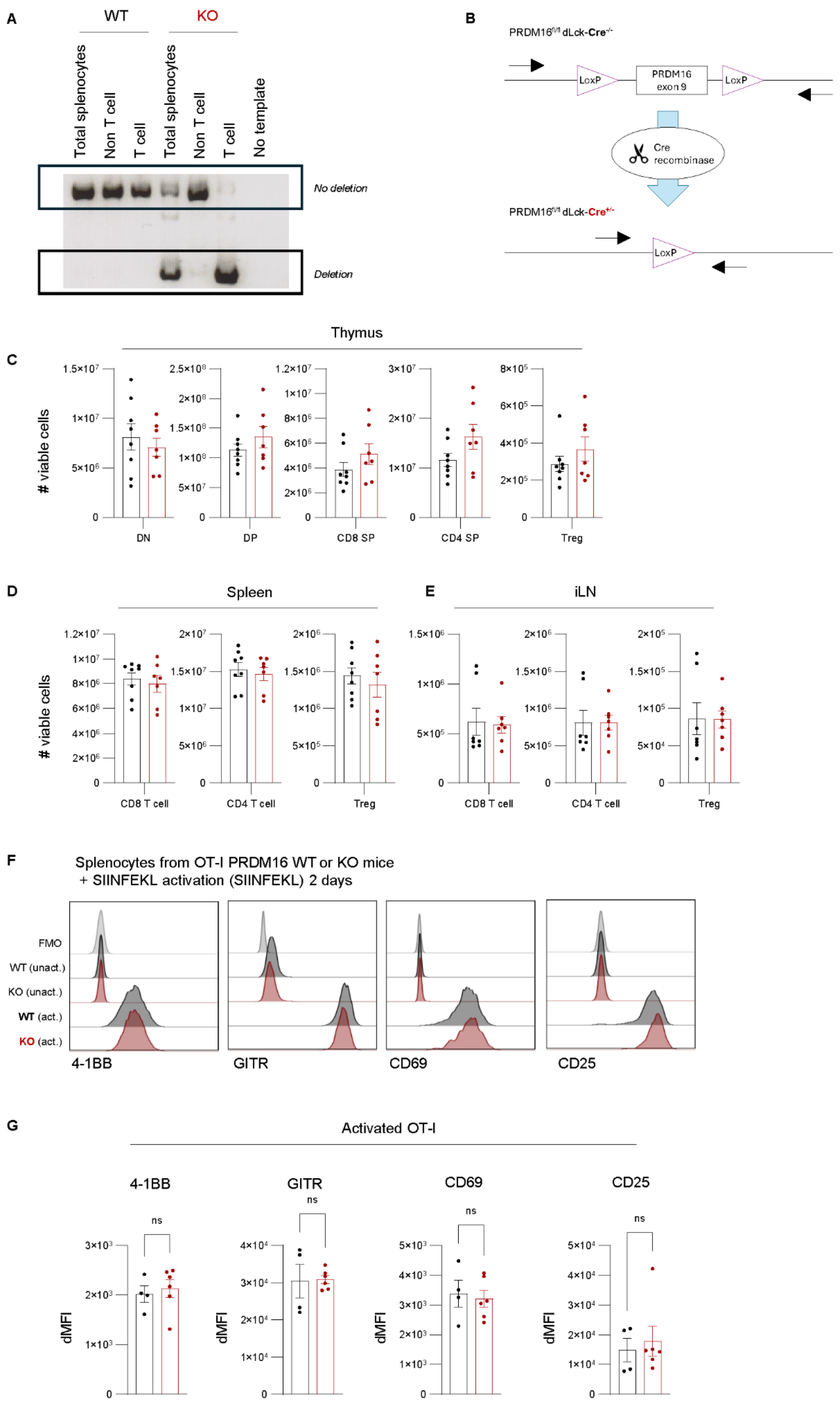


**Supplemental Figure 2. T cell-specific deletion of *Prdm16* does not affect T cell numbers in the steady state nor initial T cell activation. (A-B)** Genomic confirmation of *Prdm16* deletion in mature T cells.

Splenocytes from *Prdm16*$^{fl/fl}$ *dLck-Cre*$^{-/-}$ (WT) and *Prdm16*$^{fl/fl}$ *dLck-Cre*$^{+/-}$ (KO) littermates were isolated. T cells (Thy1.2$^{+}$CD3$^{+}$) and non-T cells (Thy1.2$^{-}$CD3$^{-}$) were FACS-sorted, and DNA was analyzed by PCR using primers that generate different product sizes depending on deletion. **(A)** shows a representative gel from two independent experiments and (**B**) shows primer orientation. **(C-E)** Cellularity of thymocytes and mature T cells in thymus (**C**), spleen (**D**), and inguinal lymph nodes (**E**) of WT and *Prdm16* KO mice. Viable cells were analyzed in the indicated subsets (n = 7 mice, two independent experiments; mean ± s.e.m.). **(F-G)** Splenocytes from OT-I *Prdm16*$^{fl/fl}$ Cre$^{-/-}$ (WT) and OT-I *Prdm16*$^{fl/fl}$ *dLck-Cre*$^{+/-}$ (KO) and mice were stimulated *in vitro* for 2 days with SIINFEKL peptide, then 4-1BB, GITR, CD69, and CD25 expression on the activated OT-I, were analyzed by flow cytometry, by gating on Thy1.2 and CD8α OT-I cells. Representative histograms are shown in **F;** quantification in **G** (n = 4-6 mice, two independent experiments; mean ± s.e.m.). Mann-Whitney U test (**C**-**E**, **G**).

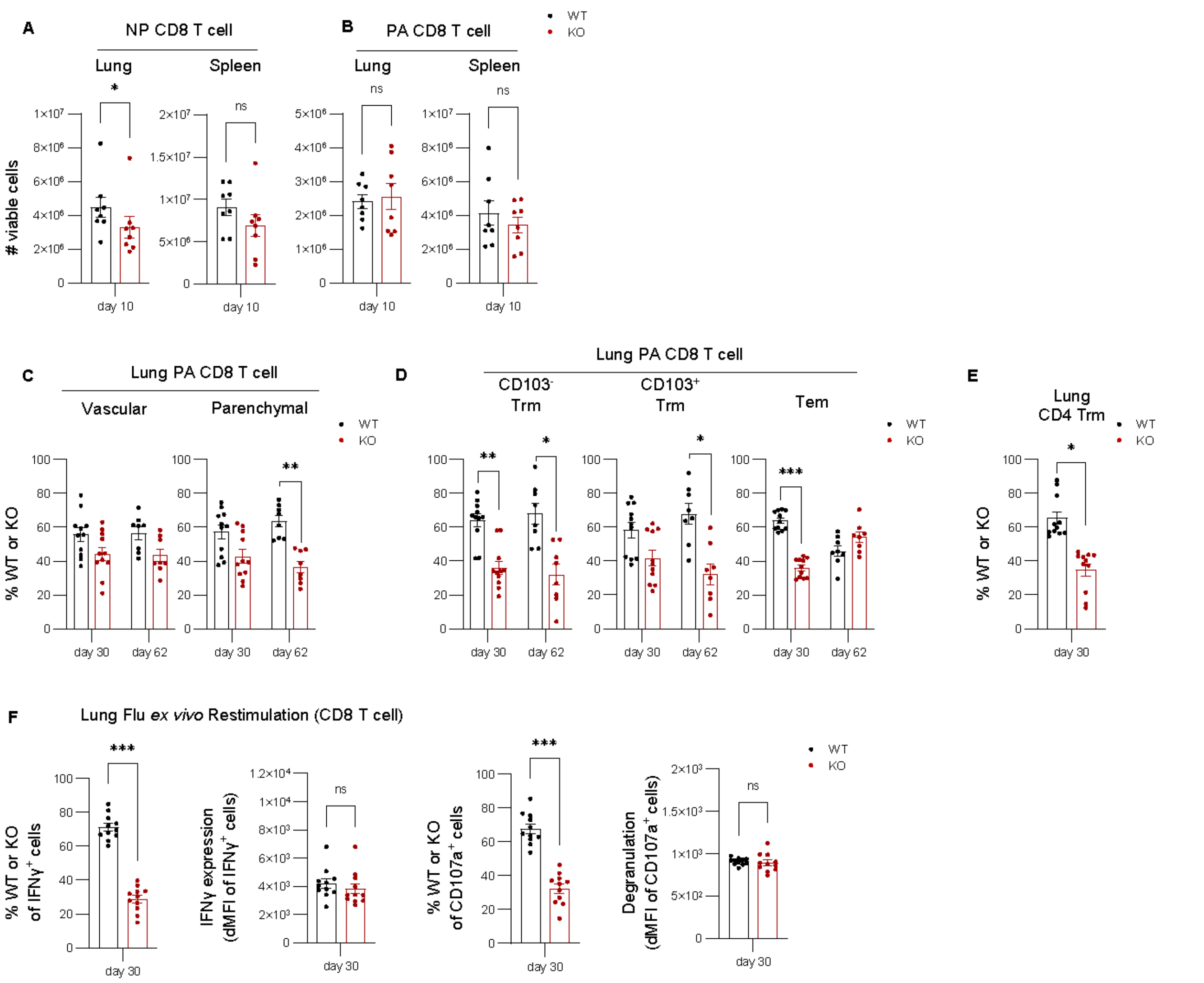


**Supplemental Figure 3. Effect of *Prdm16* deletion on polyclonal T cell responses to IAV.** *Prdm16$^{fl/fl}$ dLck-Cre$^{-/-}$* (WT) and *Prdm16$^{fl/fl}$ dLck-Cre$^{+/-}$* (KO) littermates were infected with influenza PR8 and analyzed at Day 10 p.i. Lung and spleen were analyzed for NP- (**A**) and PA- (**B**) specific CD8 T cells using MHC tetramers. Data are pooled from two independent experiments with a total of 8 mice (mean ± s.e.m.). Mann-Whitney U test (**A,B**); *p < 0.05**).** **C-E** WT: Prdm16 KO bone marrow chimeric mice were generated and infected with influenza X31 as described in Fig. 2. **(C)** Frequency of WT and *Prdm16* KO cells in lung vascular and parenchymal compartments at Day 30 and Day 62 p.i. **(D).** Frequency of WT and *Prdm16* KO cells in Trm and Tem subsets within lung parenchyma at Day 30 and Day 62 p.i. **(E)** Frequency of WT and *Prdm16* KO $CD69^{+}CD11a^{high}$ CD4 Trm cells within lung parenchyma at Day 30. **(F)** Lung homogenates were restimulated *ex vivo* with live influenza PR8 for 18 hours, with BD GolgiStop added during the final 6 hours. Frequency of WT and *Prdm16* KO cells producing IFNγ or degranulating ($CD107a^{+}$) during *ex vivo* stimulation at Day 30 p.i. is shown. WT or *Prdm16* KO frequencies p.i. were normalized to the pre-infection WT/KO ratio in blood CD8 T cells from the same mouse as described in the methods. Data are pooled from 8 to 11 individual chimeric mice in 2 to 3 independent experiments. Wilcoxon test (**C-F**, comparing pre- vs. post infection); mean ± s.e.m. ***p < 0.001, **p < 0.01, *p < 0.05

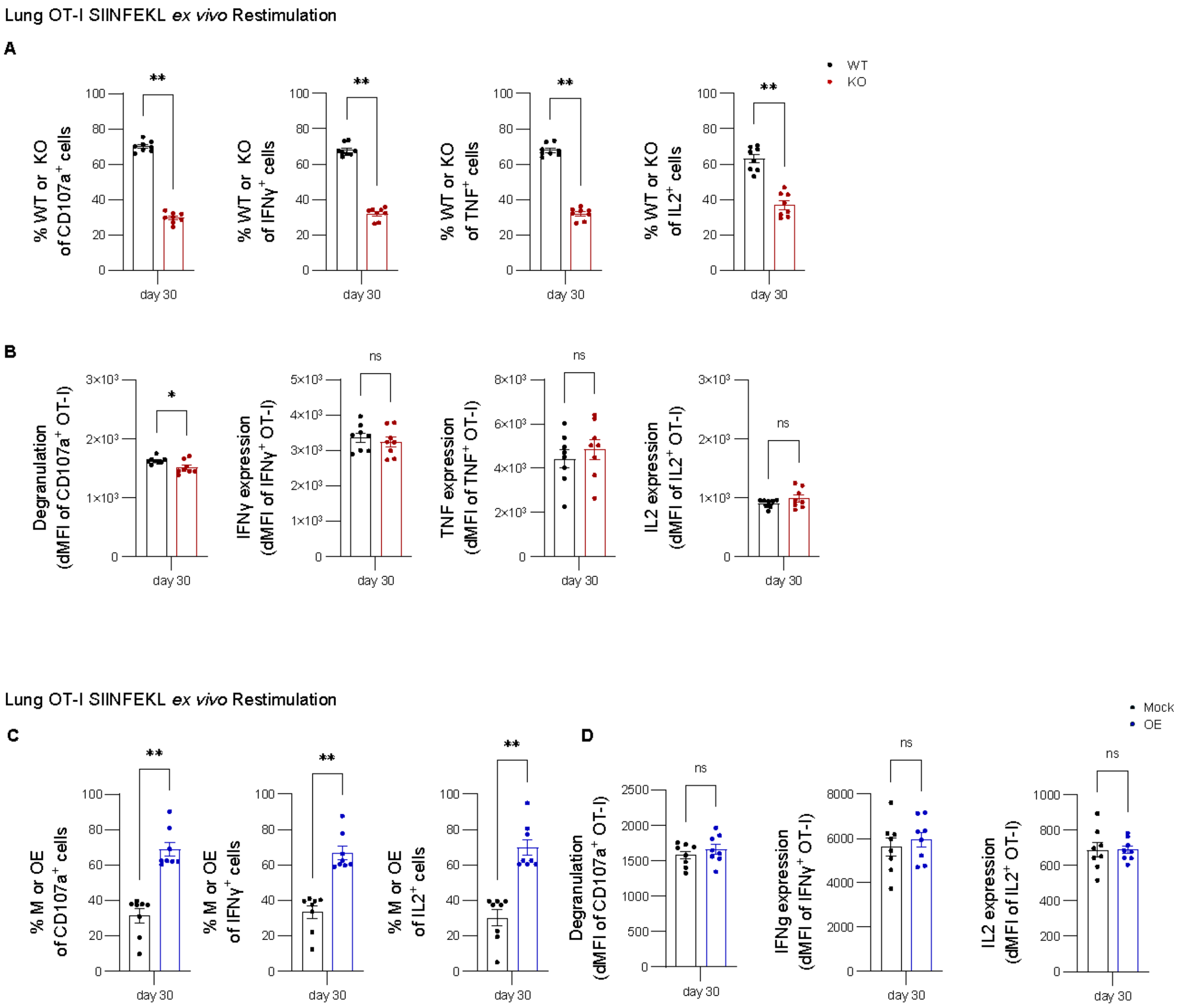


**Supplemental Figure 4. Analysis of CD8 T cell responses in *Prdm16* KO or *Prdm16*-overexpressing OT-I cells**. Lung homogenates were restimulated *ex vivo* with SIINFEKL peptide for 6 hours in the presence of BD GolgiStop from recipient mice that had received equal numbers of congenically marked *Prdm16*$^{+/+}$ *dLck-Cre*$^{+/-}$ (WT) and *Prdm16*$^{fl/fl}$ *dLck-Cre*$^{+/-}$ (KO) OT-I cells (**A-B**) or Mock-transduced (Mock; M) and *Prdm16*-overexpressing (OE) OT-I cells (**C-D**). WT or KO frequency (**A**) and dMFI (**B**) are shown for CD107a$^{+}$, IFNγ$^{+}$, TNFα$^{+}$, and IL-2$^{+}$ cells at Day 30 p.i. Mock or OE frequency (**C**) and dMFI (**D**) are shown in CD107a$^{+}$, IFNγ$^{+}$, and IL-2$^{+}$ cells at Day 30 p.i. Data are pooled from two independent experiments (8 mice). Wilcoxon test; mean ± s.e.m. $**p < 0.01$; $*p < 0.05$.

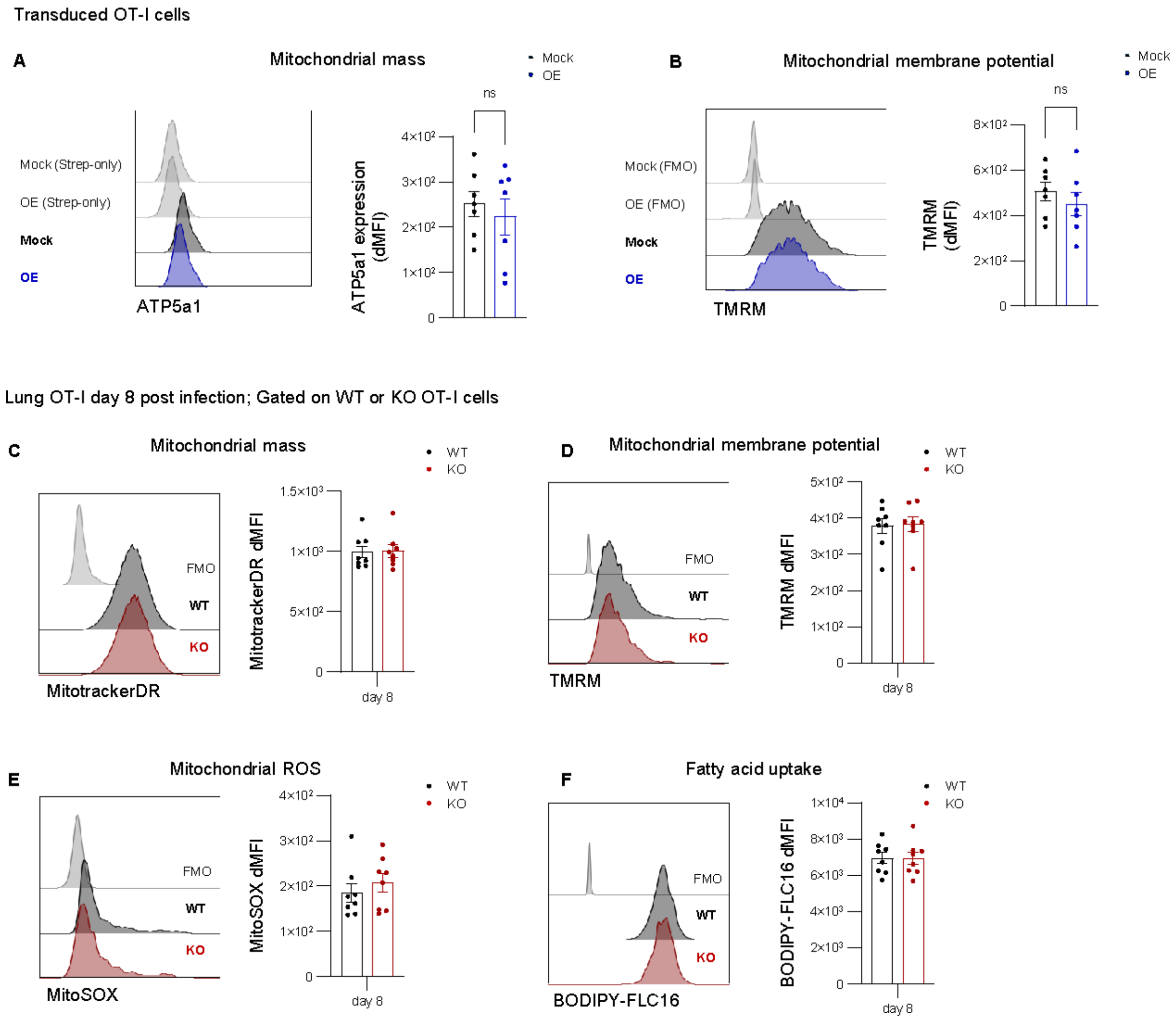


**Supplemental Figure 5. Analysis of mitochondrial state in *Prdm16* KO or *Prdm16*-overexpressing OT-I cells**. **(A-B)** Mock-transduced (Mock) or *Prdm16*-overexpressing (OE) OT-I cells were stained with anti-ATP5a1 antibody (**A**) or TMRM (**B**) to assess mitochondrial mass and membrane potential, respectively. Data are pooled from two independent experiments (7-8 mice). **(C-F)** Adoptively transferred *Prdm16*$^{+/+}$ *dLck-Cre*$^{+/-}$ (WT) and *Prdm16*$^{fl/fl}$ *dLck-Cre*$^{+/-}$ (KO) from the lung at Day 8 p.i. were stained with Mitotracker Deep Red (MitotrackerDR) **(C),** TMRM (**D**), MitoSOX (**E**), or BODIPY-FLC16 (**F**) to assess mitochondrial mass, membrane potential, ROS, and fatty acid uptake, respectively. Data are pooled from two independent experiments (total of 8 mice). Representative flow cytometry histograms and dMFI summary are shown. Statistical method: Wilcoxon test; mean ± s.e.m.

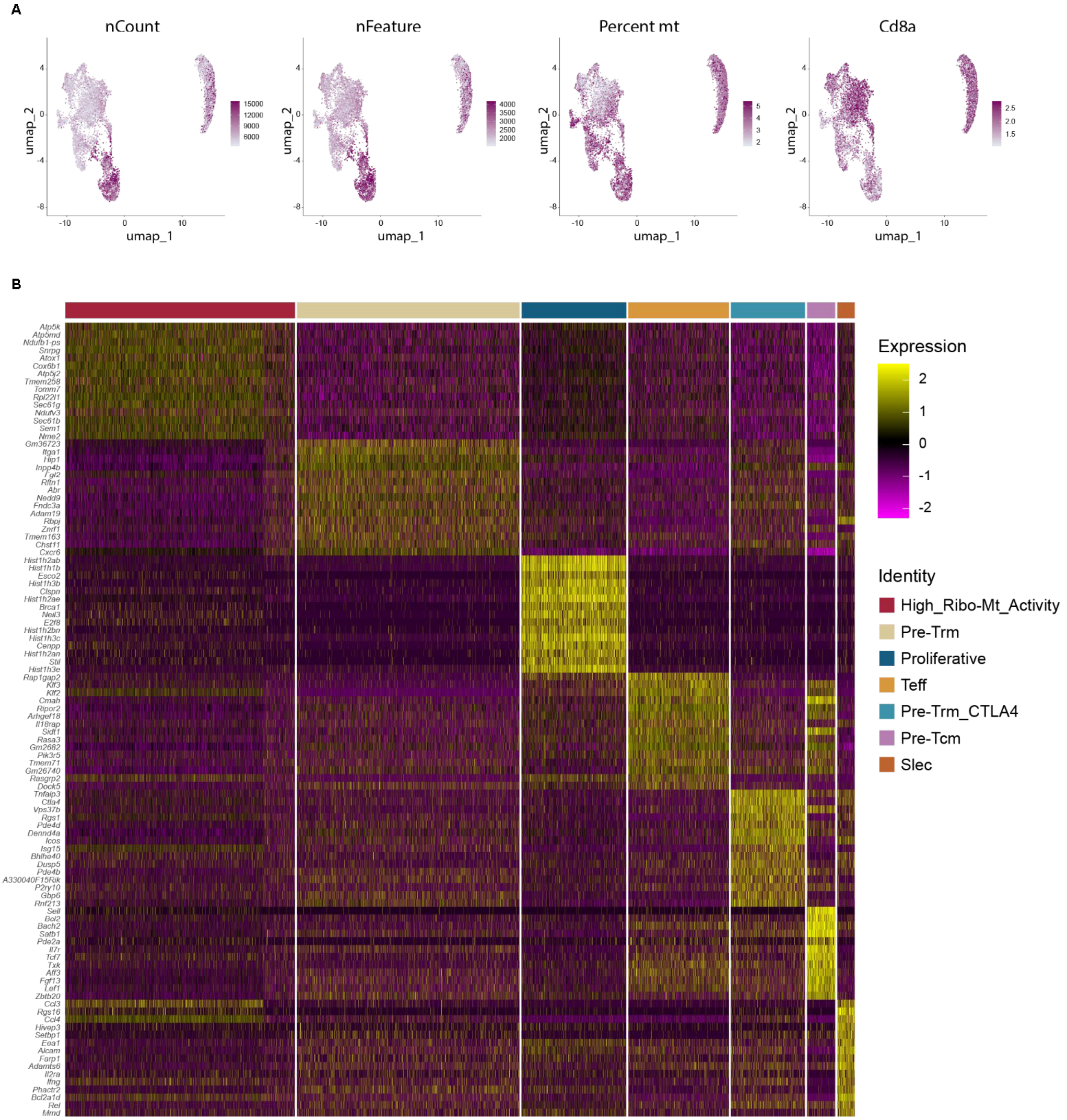


**Supplemental Figure 6. snRNA-Seq profile of *Prdm16* WT and KO CD8 T cells.** Lethally irradiated mice were reconstituted with CD45.1 (WT) and CD45.2 *Prdm16$^{fl/fl}$ dLck-Cre*$^{+/-}$ (KO) bone marrow (1:1 ratio), then infected with influenza A/X31 as described above. At day 8 post infection, CD8$^{+}$CD44$^{high}$ CD45.1 and CD45.2 cells were separately FACS-sorted from the lungs and analyzed by snRNA-seq. **(A)** UMAP showing QC features. **(B)** Heatmap of top markers for each cluster.
